# Candidate Denisovan fossils identified through gene regulatory phenotyping

**DOI:** 10.1101/2024.04.18.590145

**Authors:** Nadav Mishol, Gadi Herzlinger, Yoel Rak, Uzy Smilanksy, Liran Carmel, David Gokhman

## Abstract

Denisovans are an extinct group of humans whose morphology is mostly unknown. The scarcity of verified Denisovan fossils makes it challenging to study their anatomy, and how well they were adapted to their environment. We previously developed a genetic phenotyping approach to gain insight into Denisovan anatomy by detecting gene regulatory changes that likely altered Denisovan skeletal morphology. Here, we scan Middle Pleistocene crania for unclassified or disputed specimens that match predicted Denisovan morphology and thus might be related to Denisovans. We found that *Harbin*, *Dali*, and *Kabwe 1* show a particularly good alignment with the Denisovan profile, with most of their phenotypes matching predicted Denisovan anatomy. We conclude that our genetic phenotyping approach could help classify unidentified specimens, and that *Harbin*, *Dali*, and *Kabwe 1* exhibit a Denisovan-like morphology and could be closely linked to the Denisovan lineage.

## 2 Introduction

Denisovans are an extinct human lineage that likely shared a common ancestor with Neanderthals 390-440 thousand years ago [1, 2, 3, 4]. The first Denisovan finding was reported in 2010 based on DNA extracted from the distal phalanx of a fifth finger [2]. Since then, ten additional specimens (as well as sediment DNA) have been attributed to Denisovans based on their DNA or protein sequences [1]. This collection includes three molars [5, 3], a long bone fragment [3], a partial parietal bone (unpublished), half a mandible with two molars [6], three undiagnostic bone fragments [7] and a rib fragment [8]. These remains revealed several aspects of Denisovan morphology, including their large molars [2], a fifth distal phalanx resembling that of anatomically modern humans (AMHs) [9], and an archaic mandibular morphology which includes a robust and relatively low and thick body, without a developed chin [6]. However, the number of confirmed Denisovan specimens is still low, hindering our ability to study Denisovan adaptations and phenotypic evolution.

The majority of confirmed Denisovan remains have been discovered within the Denisova Cave in Siberia. Nevertheless, a growing body of evidence indicates that their geographical presence extended further to the east and south. This evidence includes the Denisovan sediment DNA and mandible found in the Tibetan plateau [10, 6], a tooth from Laos assumed to be Denisovan based on dental morphology [11], a mandible from Taiwan [12] exhibiting similarities in dental and mandibular morphology to Denisovans, and admixture with populations currently living in East Asia, Southeast Asia, South Asia, and Oceania [1]. Thus, it is likely that Denisovans inhabited an extensive geographical range.

Meanwhile, many hominin specimens dated to the Middle and Upper Pleistocene remain poorly classified. The fossil record of these periods is often controversial, and the ability to genetically categorize these remains is limited [13]. Consequently, certain specimens have been classified as new provisional lineages (e.g., H. *cepranensis* [14, 15], H. *bodoensis* [16], H. *mabaensis* and H. *daliensis* [17]), while others have been grouped together into broad taxonomic groups such as H. *heidelbergensis* [18], despite their high variability [19]. The difficulty in defining and classifying fossils from this period is often referred to as “The Muddle in the Middle” [20].

Importantly, many of these debated specimens have been found in East and Southeast Asia, i.e., in the likely habitat of Denisovans. Notable cranial examples include *Harbin* [21], *Dali* [22], *Jinniushan* [23], *Xuchang 1* [24], *Xujiayao* [25], *Hualongodng* [26] and *Penghu* [12] (for a recent review see [27]). Some researchers advocate for their classification as independent species [21, 13], while others have suggested that they are eastern representatives of H. *heidelbergensis* [18], or local variants of archaic H. *sapiens* [28, 17]. As academic debate continued, their taxonomy remained unclear, in what was described as a “taxonomic limbo” [17]. Following the sequencing of the Denisovan genome, some researchers proposed that some of these specimens might belong to Denisovans [29, 1]. Nonetheless, because Denisovans are a lineage defined through their genetics, but these specimens lack genetic or proteomic data, testing whether they belong to Denisovans requires bridging the gap between genetics and morphology. This can be achieved by extracting phenotypic information from the Denisovan genome and comparing it against candidate specimens.

Gene regulatory differences are a key driver of phenotypic evolution, and can be very informative of phenotypic changes between modern and archaic humans [30, 31, 32, 33]. We have previously developed a method that utilizes gene regulatory data to compare two individuals and discern which one has the higher phenotypic value (e.g., taller stature) [34, 35]. This method is based on two key conjectures: (i) substantial alterations to gene regulation are likely to have resulted in phenotypic changes, and (ii) the direction of phenotypic change following gene down-regulation is likely the same as the direction of phenotypic change in cases of gene loss-of-function [34]. To detect gene regulatory changes between the Denisovan, Neanderthal, and modern human lineages, we leveraged our previously published maps of a key regultory mark of the genome - DNA methylation [36, 37]. By linking observations of gene silencing with the reported phenotypic consequences of gene loss-of-function, we constructed a method to genetically infer morphological profiles. We first tested this approach by reconstructing a profile of Neanderthal and chimpanzee anatomy and compared them with their known morphology. We found that this method has a prediction accuracy of over 85%. Then, we applied it to the Denisovan lineage, providing 32 cranial phenotypes that likely separated it from Neanderthals, modern humans, or both.

Importantly, the phenotypic predictions that comprise the profile are qualitative rather than quantitative in the sense that they provided the direction, but not the extent, of the change. For instance, while the profile suggests that Denisovans likely exhibited a larger biparietal breadth compared to both Neanderthals and modern humans, the exact magnitude of this difference cannot be determined [34]. However, unlike trying to quantitatively predict phenotypes, predicting the direction of phenotypic difference between two individuals can reach high accuracy [35].

Here, we use the reconstructed Denisovan profile to scan the Middle Pleistocene fossil record for crania whose morphology matches the profile and thus might showcase a Denisovan-like morphology, or share close phylogenetic links with Denisovans. We identify several such specimens, including *Harbin* and *Dali* from East Asia, and surprisingly also *Kabwe 1* from Africa.

## 3 Methods

### 3.1 Selection of specimens

The majority of cranial measurements were directly taken from, or based on, the dataset provided by Ni *et al.* [21] in Morphobank [38] (project # 3385). The dataset was downloaded on 07/09/2022. Specimens in this dataset were excluded from the analysis if they:

1. Lack a cranium.
2. Pre-date the Neanderthal-Denisovan split (390-440 kya) [1].
3. Belong to a sub-adult.
4. Have fewer than five testable predictions.

This resulted in a total of ten test subjects (Figure 1). Although genetic and archaeological evidence suggests that Denisovans primarily inhabited Eastern Eurasia, we chose not to restrict our search to this region for two key reasons. First, Denisovans are already known to have occupied a wide range of geographical areas, from the Altai Mountains to Laos. This suggests the possibility of their presence in additional, as-yet-undiscovered regions. Second, the current genetic evidence for the Denisovan habitat is largely based on introgression patterns in modern populations, which primarily reflect Denisovan distribution during the Upper Pleistocene, after anatomically modern humans (AMHs) left Africa. The Denisovan habitat during the Middle Pleistocene, however, may have been significantly different. Additionally, including non-Asian specimens—despite their lower likelihood of being Denisovans—could provide insights into their evolutionary proximity to Denisovans. Beside the group of test subjects, we included 3 reference groups: (1) 20 H. *erectus*, (2) 18 H. *sapiens*, and (3) 15 Neanderthal crania (Supplementary Table 1).

**Figure 1:**
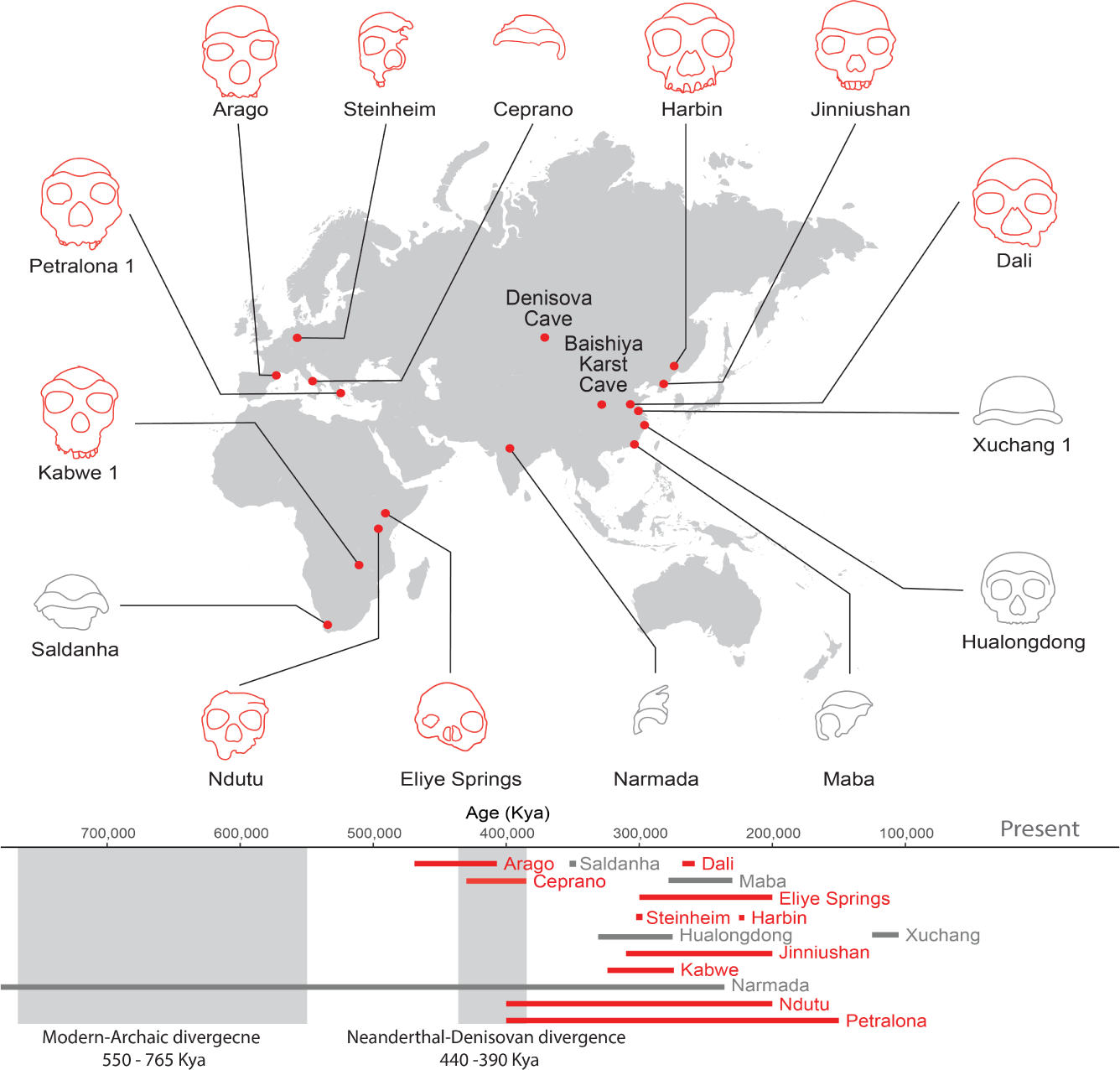
Candidate specimens tested against the Denisovan anatomical profile. In red are specimens that were included as test subjects. In grey are crania with fewer than five testable phenotypes, or that belong to sub-adults. Sites with confirmed Denisovan findings are highlighted - Denisova Cave where Denisovans were first identified, and Baishiya Karst Cave where *Xiahe 1* was found. Below are the estimated time ranges of the candidate specimens. Crania are not shown to scale.

We acknowledge that H. *floresiensis* and H. *luzonensis* share temporal and potential geographic overlap with Denisovans [39, 40]. However, they were not included in this study because they were absent from the Morphobank dataset. Despite inhabiting Southeast Asia, their overall morphology and smaller dimensions strongly suggest that they represent distinct non-Denisovan hominin lineages. Similarly, we did not include H. *naledi* specimens in this study due to their established taxonomy, based on their distinct morphology, which places them on a separate hominin lineage from Denisovans [25, 21]. H. *naledi* was not used as a reference group in this study, as no available H. *naledi* specimens were avaialble in the dataset.

### 3.2 Metric Cranial Measures

Predicted Denisovan phenotypes were taken from our original reconstruction [34]. Among these predictions, we used only the ones which could be matched against an available continuous measurement in the available dataset [21] (Supplementary Table 2)

Each predicted phenotype was associated with a continuous measurement (Supplementary Table 2) based on its description in the Human Phenotype Ontology (HPO) database [41] and other sources of facial morphology (primarily [42]). For three predictions that did not have a corresponding continuous measurement (calvarial curvature, forehead height, and glabellar protrusion), we generated a fitting measurement using available images rotated to the Frankfurt planes from Ni *et al*. [21] (see below). Overall, this resulted in ten features, representing 14 predictions of directional phenotypic differences (ten between Denisovans and AMHs and four between Denisovans and Neanderthals).

Ni *et al*. (2021) [21] provided two matrices in Morphobank: raw measurements (v1.0) and normalized measurements (v1.1). The normalized measurements were generated by dividing the raw measurements by the third power of the cranial capacity of the respective specimen to account for the effect of body size. We decided to use the raw data for several reasons: (i) the normalization method used by Ni *et al* [21]. could create potential biases between the measurements of the neurocranium versus the viscerocranium. Ideally, the size would be calculated based on the centroid of the cranium 3D model, however these values were not provided; (ii) most cranial phenotypes in the original reconstruction suggested higher values in Denisovans, possibly resulting in an overall increase in cranial size, driven by many gene regulatory changes. Such cranial expansion could be missed using a normalized dataset; (iii) The original reconstruction [34] was based on morphological characteristics that differentiate Neanderthals from AMHs, which are most often described in absolute, rather than relative, terms (e.g., when Neanderthals are described as having short extremities, it usually indicates a shorter absolute length of the extremity compared to that of AMHs) [43, 41] and; (iv) the estimation of accuracy was based on these absolute values and reached a high accuracy [34].

It should be noted that the Denisovan phenotypic reconstruction only used phenotypes where the prediction went through several quality control steps that assured high accuracy. One of these steps was to exclude from the list of Denisovan predictions in cases where the same prediction in Neanderthals was not confirmed by Neanderthal fossils. In other words, these are cases where the observed DNA methylation patterns do not accurately predict the direction of phenotypic difference [34]. In this respect, we acknowledge a mistake in our original reconstruction, where we predicted Denisovans to have thicker enamel, failing to realize that our prediction of thicker enamel in Neanderthals was wrong in the first place [44]. Therefore, as in other phenotypes where methylation was shown not to be predictive of morphology, we have removed this phenotype from the list of predictions used in the current work.

Below, we provide a detailed description of the predictions and their corresponding measurements (see also Supplementary Table 2).

#### 3.2.1 Palate breadth

The Denisovan palate was predicted to be wider than that of AMHs based on the *Narrow palate* phenotype [HP:0000189]. This phenotype is described as decreased palatal width. Here, we measured it using maxilloalveolar breadth (MAB), defined as the greatest breadth across the alveolar border, perpendicular to the medial plane [21].

#### 3.2.2 Facial breadth

The Denisovan face was predicted to be wider than that of AMHs and narrower than that of Neanderthals based on the *Small face* phenotype [HP:0000274] and the *Narrow face* phenotype [HP:0000275], which is hierarchically embedded within it. Here, we measured these phenotypes using bizygomatic breadth (ZYB, zy-zy), similarly to previous measurements of the breadth of the upper face [42].

#### 3.2.3 Facial height

The Denisovan face was predicted to be longer than that of AMHs based on the *Short face* phenotype [HP:0011219], which is hierarchically embedded within the *Small face* phenotype [HP:0000274]. This phenotype can be measured as the vertical distance between the nasion to the gnathion (the inferior border of the mandible) [42]. However, as the test subjects lacked a mandible, this measurement was unavailable. Instead, we used upper facial height (NPH), defined as the vertical distance from the nasion to the prosthion.

#### 3.2.4 Biparietal breadth

The Denisovan parietal bones were predicted to be more laterally expanded than those of both AMHs and Neanderthals based on the *biparietal narrowing* phenotype [HP:0004422]. We measured this using maximum biparietal breadth [21].

#### 3.2.5 Cranial base area

Denisovans were predicted to have a larger cranial base area than that of AMHs based on the *decreased cranial base ossification* phenotype [HP:0005451]. The cranial base is the supporting bony structure behind the midface (e.g., [45]). To estimate cranial base size, we approximated it by the area of an ellipse with one axis measured by the basion-nasion length (BNL) and the other by the biauricular breadth (AUB). Therefore, *A*_cranial base_ = *π ×* AUB *×* BNL, with both measurements taken from [21].

#### 3.2.6 Dental arch length

Denisovans are predicted to have less crowded dental arches than those of both AMHs and Neanderthals, based on the *dental crowding* phenotype [HP:0000678], described as an inadequate arch length for tooth size. Reduced dental crowding can be a result of three causes: (i) increased length of the dental arch; (ii) reduced number of teeth; or (iii) smaller teeth. The dental formula of hominins is highly consistent [46], making the second cause unlikely. The third cause is also unlikely, as the confirmed Denisovan teeth are much larger than in other *Homo* lineages [2]. We therefore concluded that the Denisovan dental arch was likely longer than than that of AMHs and Neanderthals. To test dental arch length in the maxilla, we used the maxilloalveolar length, defined as the greatest length of the alveolar process of maxilla [21].

#### 3.2.7 Facial protrusion

Denisovans are predicted to have more protruding faces than those of AMHs, but less than those of Neanderthals, based on the *flat face* phenotype [HP:0012368]. The convexity (or concavity) of the face is examined in lateral view [42]. To measure facial protrusion, we used the nasion angle (NAA) and prosthion angle (PRA). Both measurements are used as measures of prognathism [21]. Since these angles were not provided in the database, we calculated them using the law of cosines and the relevant facial measurements that were provided (NPH, BNL, BPL). The two measurements (PRA, NAA) were combined into a single value - the protrusion index, defined as the first principal component. This component explained 94% of the variability in the two measurements. We chose PCA over linear regression, as the latter assumes univariate errors (only in measuring the dependent variable), whereas in our case both variables are associated with errors.

### 3.3 Measures based on cranial images

Several measurements could not be directly matched to a suitable continuous variable from the available dataset [21]. We opted to use continuous measurements rather than the available discrete measurements for two key reasons: (1) our analysis relies on determining a central tendency, which is best represented on a continuous scale, and (2) the discrete classification of continuous phenotypes is often subjective. Instead, we used the available images from Ni *et al.* [21] to generate the continuous measurements.

The procedures for calculating calvarial flatness, forehead height, and glabellar protrusion were applied to images of the crania [21], assumed to be consistently positioned in a standard anatomical position following the Frankfurt planes. Images were provided in PNG or JPEG format. All images underwent a vectorization procedure, converting each non-white pixel into (*x, y*) coordinates. Pixels were considered white if the magnitude of the difference vector between their RGB values and standard white (255, 255, 255) was smaller than 20. The skull boundary was extracted using the Matlab alpha shape boundary algorithm with a 0.9 shrink factor. This provided an ordered list of points (*x_i_, y_i_*)*, i* = 1 *... N* on the closed curve forming the outer boundary of the cranium.

This closed curve is then expressed in terms of a series of Fourier coefficients [47]. As the cranium boundary is continuous, the coordinates of each vertex in each of the two dimensions can be thought of as a unique function of the arc length denoted as *s*, with 0 *≤ s ≤ L*, where 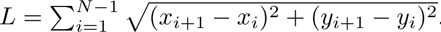. As *N* must be an even number for the subsequent procedure, in case it is odd, one vertex is removed. For convenience, the vertices are shifted by interpolation along the curve so that the distance between each two points will be constant. A discrete Fourier series is then fitted onto the points 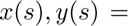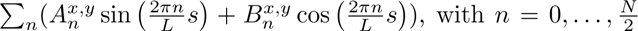 for each dimension (*x, y*). The fitting is performed by solving a system of linear equations through matrix inversion. This provides a list of Fourier coefficients (*A_n_*,*B_n_*) for each of the two dimensions, which allows the expression of the coordinates of the curve for each arc-length value.

The maximal resolution in which the curve can be expressed equals to the shortest wavelength of the Fourier series 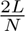. However, for some applications, for example, finding prominent low-level features such as locations of curvature peaks, it is sometimes useful to reduce the curve’s resolution. This is achieved by introducing a smoothing factor [47] that provides a weight to each term in the series 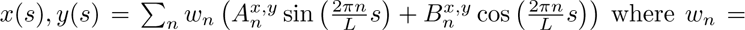 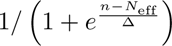. *N*_eff_ is a constant value chosen to reflect the number of terms in the series to be considered, as *w_n_ ≈* 1 for *n < N*_eff_ and *w_n_ →* 0 when *n > N*_eff_. Δ reflects an interval over which this transition gradually occurs.

The Fourier series, which enables the expression of the (*x, y*) coordinates as a function of arc-length *s* can be differentiated [47]. Its first derivative is used to express the tangent angle as a function of the arc-length 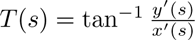. Its second derivative is used to express the curvature as a function of the arc-length 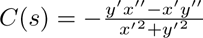.

#### 3.3.1 Calvarial curvature

Denisovans were predicted to have a calvarium (top of the cranium) that was flatter in lateral view than that of AMHs, based on the *oxycephaly* phenotype [HP:0000263]. Here, we measured calvarial curvature using a lateral image of the cranium. The anterior boundary of the cranial top was set to be the supraorbital sulcus, defined as a local negative minimum of the curvature function. The posterior boundary was set to be the mirror-reflected projection of the supraorbital sulcus on the posterior part of the cranium. This point could have been identified automatically from the curvature function. However, many crania were fragmented in different ways, which prevented an anatomically consistent choice. To avoid such errors, a semi-automatic approach was used in which we selected the correct critical points out of several potential points within a graphical user interface. The potential points are all those for which there is a local negative minimum in the curvature function and whose *x* coordinate is positive, as the critical point is concave and anteriorly positioned. The identification of the local minima is done with *N_eff_* = 45 and a buffer of 3% of the length of the clavarium around the minima, allowing the accurate identification of low-level features while filtering out points generated due to noise.

Once the critical point is selected, the arc length values corresponding to the superior cranial curve segment are plugged into the function to calculate 300 equidistant points on the curve under low smoothing with *N_eff_* equal to 80% of the number of terms in the Fourier series. These points are then scaled to centroid size 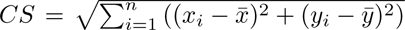 corresponding to the mean of all points on the curve. The scaled curve segment points are then fitted with the function 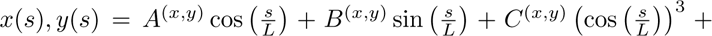 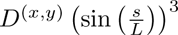 by solving an overdetermined system of linear equations. This function was chosen due to the balance it provided between compactness and error rates, allowing the expression of the general morphological aspect of the calvarium while smoothing local irregularities that may result from post-depositional damage. Similarly to the Fourier series, this function too is differentiated twice to calculate the tangent and curvature as a function of arc length expressing the angle between the tangent to the curve and its curvature.

The flatness value CF was calculated as the integral of the curvature’s absolute value over the curve segment representing the calvarium. This provides cranial flatness value of 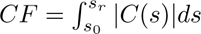 with *s_r_* being the critical point’s posterior reflection, such that lower values reflect a flatter segment (Supplementary Figure 1 a-b).

Whenever possible, the flatness value CF was calculated for both lateral right and later left images. This measure could not be computed for crania in which neither of the lateral images was intact (namely, the calvarial outline is complete and continuous). In crania where only one side was intact, only that side was used. In crania where both sides were intact, their average value was taken. We also compared the values produced from each side to examine the effectiveness of this method (Supplementary Figure 2); The error was calculated by Left-Right. One specimen (Peking 12) with outlying error values (Error *>* 2 standard deviations) was removed from further analysis. Overall, data that was available for both sides showed high Pearson correlation between them (*r* = 0.93, *p* = 2.2 *·* 10^−16^) (Supplementary Figure 2), indicating high consistency.

It should be noted that the lateral images used here were previously used for a discrete evaluation of a similar measure by Ni *et al.* [21], indicating their fit for this sort of analysis. However, we preferred to generate the measurements for forehead height and calvarial curvature ourselves, instead of using the existing discrete observations, for two main reasons: First, our methods to quantitate predictions fit better to a continuous data (see below). Second, our measurements are well-defined and therefore form an objective quantitative measure.

Our metric for calvarial curvature serves as a quantitative refinement of a similar discrete estimation in Ni *et al.* [21], who looked at the convexity along the sagittal profile of the frontal bone between the supratoral sulcus and the bregma [Discrete phenomic character #419] [21]. Then, they classified the lateral images as either *flat*, *slightly convex* and *strongly convex*. We showed that our continuous measurement is well compatible with the discrete categorization of Ni *et al.* [21] using the Kruskal-Wallis rank sum test (Supplementary Figure 3). This test was conducted in R using the kruskal.test function. The results revealed a statistically significant difference between the three groups (*P <* 8.98 *×* 10^−7^). Post-hoc analysis (Dunn test, using dunnTest() function [48] in the FSA package [49] in R) revealed that the means of all groups significantly differed from one another (*strongly convex* -*flat* adj. *P* = 4.19 *×* 10^−7^,*slightly convex* -*flat* adj. *P* = 1.41 *×* 10^−3^, *strongly convex* -*slightly convex* adj. *P* = 9.87 *×* 10^−3^).

#### 3.3.2 Forehead height

Denisovans were predicted to have a lower forehead than that of AMHs based on the *high forehead* phenotype [HP:0000348]. Similarly to the calculation of calvarial curvature, we used lateral images of the crania. A series of 300 equidistant points on the calvarium curve were calculated using the Fourier coefficients under a smoothing of *N_eff_* = 14. Then, the tangent function was used to identify the cranial vertex *s_v_*, defined as arg min*_s∈_*_[*s*_0__ *_,s_r__* _]_ *|T* (*s*)|. It was also used to locate the forehead point *s_fr_*, which was arbitrarily defined as arg 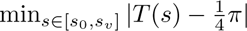. This allowed the calculation of the forehead height *FH* = *y*(*s_fr_*) *− y*(*s*_0_) with *s*_0_ being the critical point.

Similarly to the values of calvarial curvature, we evaluated the forehead height method by comparing the right and left lateral images (Supplementary Figure 4). Two specimens with outlying differences between the two sides were removed (Ngandong 7 and Steinheim). Then, the estimates drawn from the two sides of each individual showed high Pearson correlation (*r* = 0.82, *P* = 2.35 *×* 10^−12^), indicating high consistency of the method.

#### 3.3.3 Glabellar curvature

Denisovans (as well as Neanderthals) were predicted to have a retracted glabellar region compared to that of AMHs. This phenotype was originally omitted from the reconstruction [34], due to its interpretation as the absolute protrusion of the glabella, together with the supraorbital torus. Since Neanderthals are known to have a more projecting glabella than that of AMHs [50, 51], this prediction was first evaluated to be incorrect. However, after re-evaluation, we concluded that *Glabellar protrusion* in HPO refers to the protrusion of the glabellar region relatively to the rest of the supraorbital torus, and not in absolute terms [42]. The relative position of the glabella affects the curvature of the immediate glabellar region when observed in anterior and superior views. Indeed, the Neanderthal glabella was previously described as more anteroinferiorly positioned compared to AMHs and African H. *erectus*, resulting in a more receding mid-sagittal supraorbital region [51], creating a double-arched shaped supraorbital torus in Neanderthals. Since the predicted phenotype aligns with this description, we have decided to treat this as a correct prediction, in contrast with the original analysis [34].

Here, we measured glabellar curvature using superior images of the cranium. For this measurement, we only used crania that had an intact supraorbital torus in the midsagittal plane. In cases where the midfacial bones were apparent in superior view, they were digitally removed. This procedure was done for the following specimens: Sima de los Huesos 5, Dmanisi 2282, Dmanisi 2700, Saccopastore 1, Shanidar 5,Bodo, ER-1805 and OH 24. The last three specimens were not included in the final analysis as test subjects or as part of the reference groups, but were nevertheless used for the following validation of the glabellar curvature estimation method (see below).

To derive a measure of glabellar curvature, we used superior view images of the crania. Here too, it was assumed that all crania were presented in a consistent anatomical position. The boundary of the cranium and its Fourier coefficients representation were extracted using the same procedure detailed above with a smoothing factor *N_eff_* = 20. Given the cranial morphology, the boundary curve centering and the assumption of consistent rotation, the arc length value of the glabella *s_gl_* was that for which *x*(*s_gl_*) *>* 0*, y*(*s_gl_*) = 0, or in other words the intersection of the curve and the *x* axis in its positive part. Thus, the glabellar region was defined as the segment of the curve ranging between 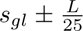 (Supplementary Figure 1 c). The value of the curvature function was calculated for *s_gl_*, as well as for another 20 equidistant points covering this section. The glabellar curvature is expressed as *C*(*s_gl_*). In cases where there was noticeable cranial asymmetry that prevented *s_gl_* from corresponding to the anatomical glabella, we selected the value of the correct point out of the 20 points for which curvature was calculated. The glabellar curvature values for all groups are presented in Supplementary Figure 1 c.

Similarly to quantitive refinement of the calvarial curvature estimation, a similar discrete metric was also used by Ni *et al.* [21], termed “Glabella concavity relative to the supraorbital tori”. This metric was based on superior view images, and included three categories: *deep*, *shallow*, and *absent*. Here too, we used the Kruskal-Wallis test to compare between the discrete metric of Ni *et al.*[21] and our continuous measure (Supplementary Figure 6). In support of the compatibility between the two measures, we found a statistically significant difference between the three groups (*P <* 3.04 *×* 10^−7^). Post-hoc analysis (Dunn test) revealed that the *absent* group is significantly different from the other groups (*absent* -*deep* adj. *P <* 1.74 *×* 10^−6^, *absent* -*shallow* adj. *P* = 3.31 *×* 10^−4^). However, the *shallow* and *deep* groups do not significantly differ from each other (adj. *P* = 0.266). Overall, these results serve as further support for the effectiveness of the method we developed.

The curves used to calculate calvarial curvature, forehead height, and glabellar curvature can be found in Supplementary Figure 7. Examples of the curves used for the reference groups can be found in Supplementary Figure 8.

### 3.4 Quantile estimation

Each single comparison was carried out by computing the percentile of a particular measurement in a certain test subject with respect to the distribution of this measurement in the reference group (either Neanderthals or AMHs). To this end, let *t* be the value of the measurement in the test subject, and let *t*_1_ *≤ t*_2_ *≤ . . . ≤ t_N_* be its values in the reference group. The percentile was computed using the following steps: (1) Outliers were removed from the reference group. Values below *Q*_1_ *−* 1.5 *×* IQR or above*Q*_3_ + 1.5 *×* IQR were considered outliers. (2) The value *t_i_*in the reference group was taken as the estimator of the (*i−*0.5)*/N* quantile, 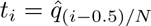. The quantile of tied values was taken as the mean of the respective quantiles (the parameter ‘ties’ was set to ‘mean’). (3) The remaining quantiles were linearly interpolated using the approxfun function in R, to create a cumulative distribution function (CDF). (4) the percentile of *t* was estimated based on the generated CDF.

### 3.5 Phenotypic distance

Based on the estimated quantile of a measurement with respect to a reference group, we defined a “phenotypic distance” that provides a normalized score for the distance of this measurement in the test subject from the reference group. This distance lies in the range (*−*1, 1), with positive values reflecting agreement with the prediction in Denisovans, and negative values reflecting disagreement. To compute this distance, quantiles should first be centered around 0 by subtracting 0.5. Then, the predicted direction of phenotypic change is accounted for by flipping the sign if the prediction was that the measurement in Denisovans was lower than in the reference group. Finally, the distance is scaled to (*−*1, 1) using multiplication by two. Let *s* be +1 if the measurement is predicted to be higher in Denisovans, and -1 if it is the other way around. Let *q* be the quantile of the measurement in a specific test subject with respect to the reference group. Then, the phenotypic distance would be

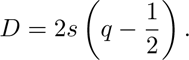

Because measurements that align with the Denisovan profile result in positive distances, specimens resembling the Denisovan profile exhibited an asymmetric distribution skewed toward positive values (e.g., Fig. 11a). Conversely, specimens with limited resemblance to the Denisovan profile tended to display values that were either evenly dispersed or skewed toward negative values (e.g., Fig. 11e). Importantly, a higher positive value does not necessarily mean a better fit to the Denisovan profile, as the directional predictions are qualitative and not quantitative, and thus, the typical Denisovan quantile is unknown, except for whether it is predicted to be greater than or lower than 0.5. However, a higher phenotypic distance does indicate increased confidence in the observed divergence between the test subject and the reference group.

### 3.6 Scoring the match between a test subject and the Denisovan profile

Let *d*_1_*, d*_2_*, . . ., d_k_* be the phenotypic distances of *k* different measurements obtained for a particular test subject. Based on these distances, we scored how closely the test subject resembles the predicted Denisovan profile using two approaches. First, we consider a measurement to be compatible with the prediction if the corresponding phenotypic distance is greater than zero. Given that under the null hypothesis we have equal chances for a positive and a negative phenotypic distance, we used a one-tailed binomial test to test whether the number of positive values is significantly greater than expected by chance. The binomial test was carried out using the ‘binom.test’ function in R with continuity correction [52, 53]. Second, we used one-tailed Wilcoxon test to check whether the median of the phenotypic distances significantly deviates from zero. This was implemented using the wilcoxsign test function from the R ‘coin’ package [54]. We used this function instead of wilcox.test function since its implementation of the test accepts tied values in the dataset. The *P* -values generated by each of the two tests were then corrected for multiple comparisons using the BenjaminiHochberg procedure [55]. The multiple correction was performed only for test subjects, as all other specimens were not hypothesized to potentially belong to Denisovans or their close relatives and therefore, cannot be considered discoveries.

The phenotypic features examined in this work are correlated, which violates the assumption of both statistical tests. Hence, we do not assign a statistical meaning to the *P* -values, but rather treat them as scores reflecting the relative similarity of each specimen to the predicted Denisovan profile (see next section for a test that accounts for the correlation between features). The *binomial score* is based on the binomial test *P* -value, while the *wilcoxon score* is based on the Wilcoxon test *P* -value. The scores were calculated from the *P* -values as

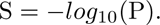

### 3.7 Permutations

A permutation test was applied to all test subjects in order to validate the fact that the measures we used in the actual analysis are significantly more informative than a random set of measures. Each permutation was generated by sampling random measures from the pool of available continuous measures in the Morphobank dataset of Ni *et al.* 2021 [21]. For consistency, the five measures generated for the current work were included in the pool as well (these include calvarial curvature, forehead height, glabellar curvature, cranial base area and facial protrusion). For each specimen, the original measurements were replaced by other randomly sampled measurements, which were then used to score the resemblance of the test subject to the predicted Denisovan profile.

To avoid introducing any biases, the permutations were designed to conserve all relevant properties of the original analysis. First, the number of testable predictions was maintained across all permutations of each test subject. Since the preservation level of each specimen differs, the number of testable predictions is not equal for all specimens. For example, the *Petralona 1* cranium is highly preserved and has 14 testable predictions. In contrast, *Ndutu* has only 9 testable predictions, due to poorer preservation.

Second, the number of measures compared to each reference group (AMHs, Neanderthals) was kept fixed for each test subject. For example, *Petralona 1* has 10 measurements that were compared to AMHs, and four that were compared to Neanderthals. These numbers were retained in all *Harbin* permutations.

Lastly, the original analysis used a predicted directionality of each measure compared to a reference group. However, this directionality is arbitrary, as it is based on the way the measure is defined. For example, a measure defined as forehead height is predicted to be smaller in Denisovans compared to AMHs. However, were we to define the measure as forehead shortness, this measure would have been predicted to be larger in Denisovans. In order to account for this when carrying out the permutations and to maintain the relationships between different measures, we redefined the directionality of each measure based on its correlation with biorbital breadth (EKB). Measures with a positive correlation with EKB (*n* = 118) were considered to have positive directionality, while measures with a negative correlation (*n* = 24) were considered to have negative directionality. Most measures were positively correlated with EKB, as expected given its correlation with overall size.

By maintaining the directional correlation between predictions, the number of measures per specimen, and the number of comparisons with each reference group, we controlled for potential biases, such as overall cranial size, correlations between phenotypes, and preservation level.

For each specimen, *N* = 1, 000 permutations were performed. A combined statistic which takes into account the Wilcoxon score and the binomial score 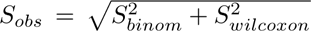 was computed and compared to permuted combined scores to produce the permutation p-value: 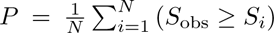. *P* -values were assigned based on the fraction of permutations in which the random match exceeds the observed one.

### 3.8 Principal Component Analysis

Since most crania used in this study are incomplete, many measurements were missing. To compute PCA, missing data had to be imputed. To this end, we used the ‘missMDA’ package [56]. This package imputes missing data by iterative PCA, which takes into account the correlation between variables and similarity between samples. Values were imputed using the function ‘imputePCA’ with default parameters: scaled data and two dimensions.

To avoid over-imputation of the data, we first filtered out measures with over 20% missing values (i.e., kept only measures where 80% or more of the specimens had data), and then specimens with more than 15% missing values. These values were selected in order to keep a maximum number of specimens while still keeping imputation low. We also computed an additional metric PCA in which we first applied the 15% specimen filtering and only then the 20% measures filtering, in order to try to retain more measures (Supplementary Figure 9).

In addition to the main PCA, which was computed using all continuous measures, we computed a second PCA strictly for non-metric measures (e.g., angles, ratio) (Supplementary Figure 10). This was done in order to test the potential effect of overall size biases on the clustering. Since there are fewer nonmetric measures in the dataset, we used a slightly more permissive threshold (30% instead of 20%) for maximum missing values per measure for this PCA. The overall imputation remains the same since the maximum missing values threshold per specimen remains the same, although fewer specimens are kept in the non-metric PCA.

In all PCAs, the analysis was based on the reference groups, and the test subjects were later projected onto this plane to produce the final plot.

### 3.9 *Xiahe 1* Mandibular analysis

For the mandibular analysis we used the mandibular data in [10]. AMH values are a weighted mean of the Early Homo sapiens, Late Homo sapiens and Asian H. sapiens groups. Neanderthal values represent the weighted mean of Asian Neanderthals and European Neanderthals.

In our previous paper [34] we reported that our reconstruction correctly predicted seven out of eight Denisovan phenotypes based on the morphological description of *Xiahe 1* [10]. Here we repeated this analysis, but this time retained only cases of directional predictions (i.e., not including cases where no difference in regulation was detected).

Four measures were examined for the comparison of the *Xiahe 1* mandible:(*i*) dental arch length (for dental arch length prediction), (*ii*) Symphesial height (for chin height prediction), (*iii*) bicanine breadth and (*iv*) bicanine breadth to Bi-M2 breadth ratio (for pointed chin prediction). For the dental arch length and pointed chin predictions we used the distance form incisors to M2 and not to M3 due to M3 being missing in *Xiahe 1*.

### 3.10 Map

The background world map used in Fig. 1 is based onthe world map by PT Northern Lights Production, used under GPLv.2 license. Available at https://mapsvg.com/.

## 4 Results

### 4.1 Denisovan specimens validate Denisovan genetic phenotyping

Morphologically informative Denisovan specimens could be used to further evaluate the accuracy of the Denisovan profile. These include *Xiahe 1* [10], the Denisova 3 distal phalanx [9], and several molars [5, 3]. Denisovans were predicted to have a longer dental arch than both AMHs (average 52.58 mm) and Neanderthals (average 54.78 mm). *Xiahe 1* has a dental arch length of 55.7 mm, compatible with the prediction in both cases. Denisovans were also predicted to havea longer symphyseal height than that of AMHs (average 32.35 mm). *Xiahe 1* has symphyseal height of 32.6 mm, again compatible with the prediction. Denisovans were also predicted to have an anterior mandible that is wider than that of AMHs and narrower than that of Neanderthals. The *xiahe 1* anterior mandible has a width (bi-canine distance) of 42.6, which is wider that of both AMHs (33.14) and Neanderthals (36.47). Next, Denisovans were predicted to have a wider anterior-to-posterior breadth ratio compared to AMHs (average 0.52). *Xiahe 1* has a wider anterior-posterior ratio of 0.57. Lastly, Denisovans were predicted to have a more protruding mandible compared to AMHs, similarly to Neanderthals. We were not able to find a direct measure for mandibular protrusion, however the authors report a mental foramen which is located under P4, more anteriorly positioned than in AMHs. Overall, six out of seven predictions were confirmed using the *Xiahe 1* mandible. This reflects an accuracy of 85%, in line with the reported accuracy of our reconstructed method [34].

The Denisova 3 distal phalanx [9] could potentially be used to test the prediction of reduced tapering of the finger from base to fingertip in Denisovans compared to AMHs. However, this comparison requires a Denisovan proximal and middle phalanges, which have not been found to date, leaving the accuracy of this prediction to be determined. Finally, despite the availability of Denisovan molars [5, 3], our predictions of Denisovan dentition relate to the timing of tooth eruption and loss, which cannot be determined using current specimens (see Metric Cranial Measures in Methods for *tooth enamel*).

### 4.2 Test subjects and measurements

Given the high accuracy of the genetic phenotyping approach, observed both in confirmed Denisovan specimens and in validating the method on Neanderthals and chimpanzees [34], we turned to scan the fossil record for specimens matching the reconstructed Denisovan profile. We started by generating a set of candidate fossils, that we denote *test subjects*. This set was done using several criteria. First, we only examined specimens whose dating partially or completely postdates the emergence of the Denisovan lineage, i.e., the estimated Denisovan-Neanderthal split (390-440 kya, [1]). Second, non-Denisovan specimens whose taxonomy is well-defined (e.g., Neanderthals, Homo *erectus*, and AMHs), were not included in the set of text subjects but rather as *reference groups*. Due to the ongoing debate about the monophyly and taxonomy of Homo *heidelbergensis* specimens ([57, 58]), as well as their phylogenetic proximity to Denisovans, which led some to suggest that some Homo *heidelbergensis* may in fact be Denisovans [29], these specimens were included among the test subjects. Third, to allow for enough power in the analysis of each specimen, and because the predicted Denisovan profile contains many cranial features, we focused on sufficiently complete crania, containing at least five testable phenotypes. Fourth, we did not include sub-adult specimens, as they could not be directly compared to the adult specimens of the reference groups. Finally, to minimize biases and to account for the possibility that the Denisovan habitat extended beyond Eastern Eurasia, we included only adult crania and did not filter out specimens based on their location. Altogether, the set of well-preserved Middle Pleistocene cranial test subjects included ten specimens (Fig. 1). These test subjects lack genetic characterization, display substantial morphological diversity, and their phylogenetic relationship to known hominin groups is uncertain or subject to debate.

The reconstructed Denisovan profile included directional predictions of 19 cranial phenotypes in which Denisovans are expected to differ from AMHs, Neanderthals, or both (Supplementary Table 2). Four of these phenotypes are non-morphometric (teeth loss timing, teeth eruption timing, skeletal maturation timing, mineralization density), rendering them unquantifiable in the test subjects. The enamel thickness prediction was dropped as well,(see Metric Cranial Measures in Methods)Out of the remaining 14 phenotypes, ten were cranial (excluding the mandible), and since none of the test subjects has a mandible, we focused on these ten phenotypes (Supplementary table 2).

Out of these ten phenotypes, seven could be matched against measures in the Morphobank dataset [21]. For the other three (calvarial curvature, forehead height, and glabellar protrusion), the Morphobank dataset has no matching information. Therefore, we developed a method to measure them from standardized images of the specimens (Fig. 2A, see Methods). The same measures (Supplementary Table 2) were then taken from 20 H. *erectus*, 15 Neanderthals, and 18 AMHs, which serve as reference groups.

**Figure 2:**
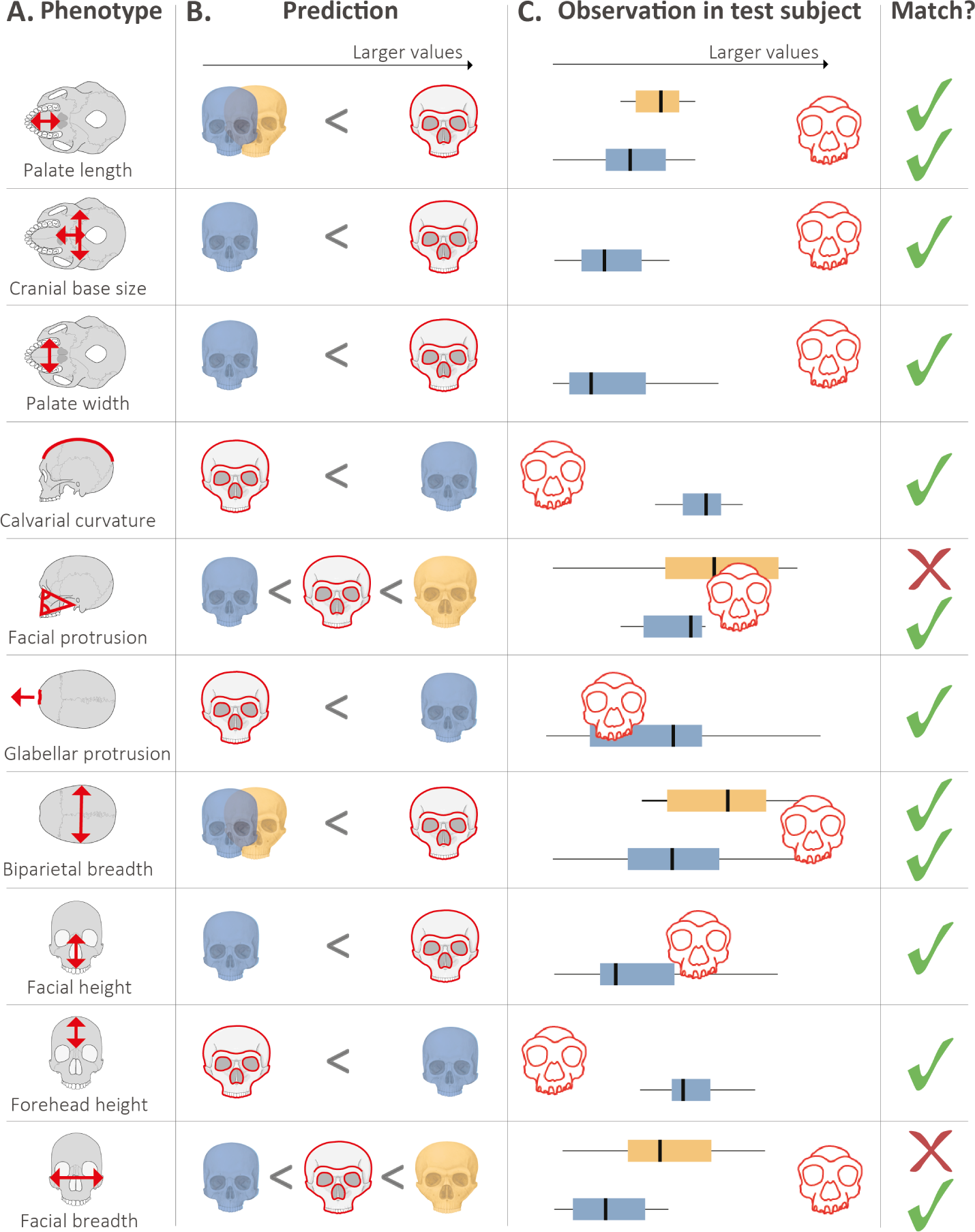
Comparing the morphology of test subjects to the predicted Denisovan morphology. **A.** Phenotypes used to evaluate the match of each test subject to the predicted Denisovan profile. **B.** Predicted relative value of each phenotype in Denisovans (red outline), compared to AMHs (blue) and Neanderthals (yellow). **C.** Boxplots showing thedistribution of each measure in AMHs (blue) and Neanderthals (yellow). The red cranium shows the value of this measure in a test subject, here demonstrated using *Harbin*. Check and cross marks show whether the *Harbin* cranium matches the predicted relative position of the phenotype in Denisovans.

In all ten phenotypes, Denisovans are predicted to differ from AMHs, and in four of them, they are also predicted to differ from Neanderthals (Fig. 2B). Thus, depending on the degree of cranial preservation, up to 14 comparisons were carried out for each test subject. In each comparison, a phenotype in the test subject was compared to the distribution of the same phenotype in a reference group – either Neanderthals or AMHs. The phenotype in the test subject was considered a match to the Denisovan profile if it fell on the side of the median of the reference group in which Denisovans are expected to fall. For instance, our profile suggested that Denisovans had a lower forehead compared to AMHs. We therefore compared forehead height in each test subject against the distribution of forehead heights in AMHs. If a test subject’s forehead height fell below the median value of forehead heights in AMHs, it was considered a match (Fig. 2C).

### 4.3 Estimating the match between test subjects and the Denisovan profile

We estimate the degree of similarity of a specimen to the reconstructed Denisovan profile using a metric we refer to as the phenotypic distance (see full description in Methods). A phenotypic distance is separately computed for each comparison of a phenotype to a reference group, and it is based on the quantile of the phenotype in the test subject compared to its distribution in the reference group. The phenotypic distance is scaled to (*−*1, 1), with positive values reflecting agreement with the predicted Denisovan profile, and negative values reflecting disagreement. It is important to note that a higher positive value does not necessarily mean a better fit to the Denisovan profile, as the directional predictions are qualitative and not quantitative, and therefore, the expected phenotypic distance of Denisovan phenotypes from the reference groups is unknown. However, higher positive phenotypic distances do indicate increased confidence in the observed divergence between the test subject and the reference group.

Because measurements that match the Denisovan profile yield positive phenotypic distances, specimens resembling the Denisovan profile exhibit a distribution of phenotypic distances that is skewed toward positive values (e.g., Fig. 11a). Conversely, specimens lacking resemblance to the Denisovan profile tend to display values that are either evenly dispersed around zero or skewed toward negative values (e.g., Fig. 11e). As control, we computed phenotypic distances for each specimen in the reference groups by treating it as a test subject, i.e., excluding it from its reference group and testing it against the reference groups.

We use the fact that positive phenotypic distances represent a match to the predicted Denisovan profile to develop two scores that measure the degree of concordance between each test subject and the predicted Denisovan profile. The first test is based on the proportion of matches of each specimen to the profile. For each specimen, we computed a *binomial score*, using the null hypothesis that that specimen is equally likely to be higher or lower than the reference group (i.e., the probability of a positive phenotypic distance is 0.5). In this analysis, higher values correspond to a greater bias towards positive values of the phenotypic distances, reflecting a better match to the Denisovan profile (see Methods).

In the second test, we accounted for the values of the phenotypic distances by testing the extent to which the median of the phenotypic distances of each specimen deviates from zero. To this end, we defined a *Wilcoxon score* based on a one-tailed Wilcoxon test for each specimen, which assigns higher values to specimens that deviated more extremely in the direction of the predicted Denisovan profile across many phenotypes (see Methods).

In summary, the binomial score primarily focuses on the number of matches, while the Wilcoxon score additionally factors in the measured value of the phenotype, giving more weight to more extreme differences between the test subject and the reference groups. Because the exact extent of phenotypic change in Denisovans is unknown, it remains uncertain which score better represents the match to the Denisovan profile. Consequently, and given the fact that the two scores are correlated (Fig 3), we utilized both to rank the specimens.

**Figure 3:**
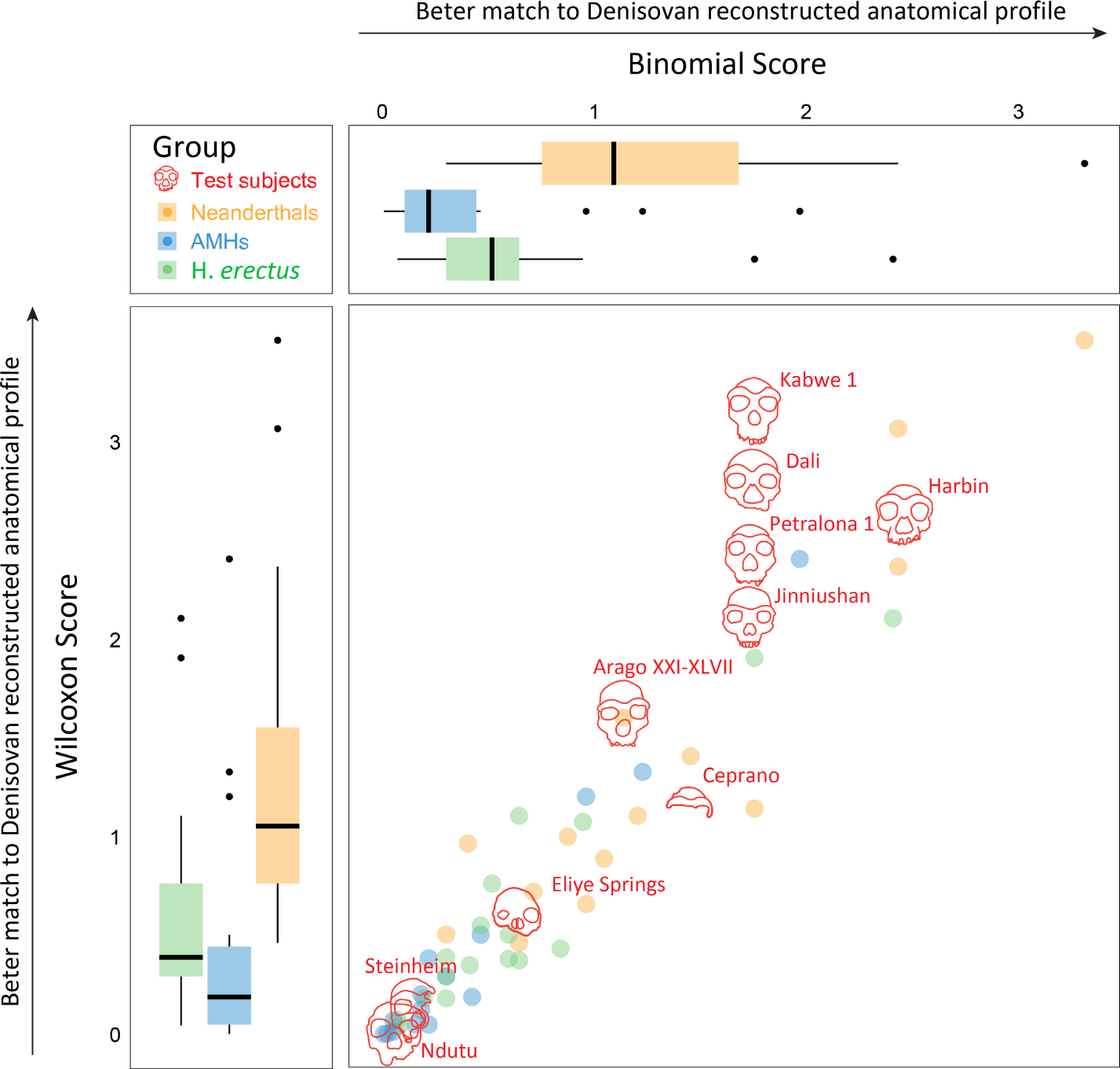
Testing various specimens against the predicted Denisovan profile. Each specimen was tested for up to 14 phenotypes predicted to distinguish Denisovans from AMHs or Neanderthals. The scatter plot shows the match of each specimen to the Denisovan profile using two scores – binomial (x-axis), and Wilcoxon (y-axis). Boxplots show the distributions of scores for the reference groups, serving as controls.

### 4.4 *Harbin*, *Dali*, and *Kabwe 1* show high concordance with the Denisovan profile

Overall, most test subjects show a higher proportion of positive phenotypic distances compared to the reference groups (Fig. 3).

In the binomial scoring, we found that the test subject showing the highest compatibility with the Denisovan profile was *Harbin*. Out of 14 phenotypes predicted to distinguish Denisovans from AMHs or Neanderthals, *Harbin* showed 12. This represents a match rate of 86% (binomial score = 2.43, see x-axis in Fig 3). This is in contrast with the substantially lower scores observed in the controls; AMH and H. *erectus* specimens show binomial scores closer to 0, reflecting match rates to the Denisovan profile that are close to the random match rate of 0.5 (0.54 for AMHs, and 0.56 for H. *erectus*). Neanderthals present higher scores, in line with their close phylogenetic proximity to Denisovans and their high predicted morphological resemblance to them [34].

In the Wilcoxon scoring, we found that the three test subjects showing the highest compatibility with the Denisovan profile were *Kabwe 1*, *Dali*, and *Harbin* (see y-axis in Fig 3). Here too, the reference groups scored relatively low, showing random match rates, whereas Neanderthals showed higher resemblance to the Denisovan profile. Overall, three specimens ranked consistently high in both scores: *Harbin*, *Dali*, and *Kabwe 1*.

If the predicted Denisovan profile holds true information about Denisovan morphology, and if a test subject indeed has Denisovan-like morphology, this should be reflected by its match scores being significantly higher than expected by chance. We implemented a permutation test in order to (*i*) validate that the measures we used to compare test subjects to the predicted Denisovan profile are more informative than randomly chosen measures, and (*ii*) estimate the accuracy of individual test subject scores. In each permutation, we randomly replaced phenotypes from the Denisovan profile with random ones from the Morphobank dataset [21], while maintaining the directional correlations between phenotypes. By doing so, we controlled for potential biases, such as overall cranial size, correlations between phenotypes, and the number of available comparisons. Then, we tested the match of each specimen to the randomly permuted profile. We repeated this 1,000 times for each specimen and assigned *P* -values by computing the proportion of permutations where the match to the permuted profile exceeded that of the match to the true profile (see Methods).

We found that most test subjects do not show a significant match to the Denisovan profile, suggesting that their morphology is not more similar to the predicted Denisovan morphology than expected by chance. However, four test subjects showed significant resemblance to the Denisovan profile: *Dali* (*P* = 0.013), *Harbin* (*P* = 0.027), *Kabwe 1* (*P* = 0.029), and *Jinniushan* (*P* = 0.039, Supplementary Table 3). Overall, we propose that *Harbin*, *Dali*, and *Kabwe 1* present unique and significant Denisovan-like morphology (see Discussion for the interpretation of this similarity).

### 4.5 High-scoring specimens exhibit morphological similarity

To explore potential morphological affinities between test subjects and examine if distinct clusters emerge, we carried out a principal component analysis (PCA). To this end, we used all available continuous cranial measures in the Morphobank dataset [21], with a maximum of 20% missing data. To this, we added the three measures generated for this study (glabellar curvature, calvarial curvature, and forehead height), leaving us with a total of 50 measures (Supplementary Table 4). To avoid over-imputation, we also filtered out specimens with more than 15% missing measurements, leaving us with a set of 50 specimens in the final PCA. The remaining missing data were imputed using the Singular Value Decomposition (SVD) method [56]. This analysis also allowed us to study *Xuchang 1* and *Narmada*, which did not have sufficient data to be compared against the Denisovan profile, but could be compared against other specimens using the 56 aforementioned measures.

The PCA exhibited a high level of separation between the reference groups H. *erectus*, Neanderthals, and AMHs (particularly between H. *erectus* and the other groups, Fig. 4). This suggests that these cranial measures are able to capture the distinct evolutionary histories of these human lineages. Next, we projected the test subjects onto the PCA plane (Fig. 4). Notably, several of them were placed close to one another and separately from the reference groups. These include *Harbin*, *Xuchang 1*, *Ceprano*, *Petralona 1*, *Kabwe 1*, *Dali*, and *Narmada*. *Jinniushan* is positioned close to Dali, between the Neanderthal and H. *erectus* populations. Other test subjects were positioned either within the other populations (e.g., *Eliye Springs*), or far from any other population (*Steinheim*). The top-right cluster includes specimens that display a high concordance with the Denisovan profile, and particularly the top three *Harbin*, *Dali*, and *Kabwe 1*. This suggests that specimens resembling the Denisovan profile form a cluster of morphologically similar specimens, not only with respect to their Denisovan-like phenotypes, but also to the rest of their cranial morphology. Furthermore, we have carried out a second PCA, using solely non-metric measures, which exhibited similar clustering patterns and morphological relations between specimens (Supplementary Figure 10). This provides further credence to their grouping in the same cluster, regardless of the overall cranial size. Interestingly, *Xuchang 1*, whose high similarity to the Denisovan profile was previously noted [34], clusters very close to *Harbin*.

**Figure 4:**
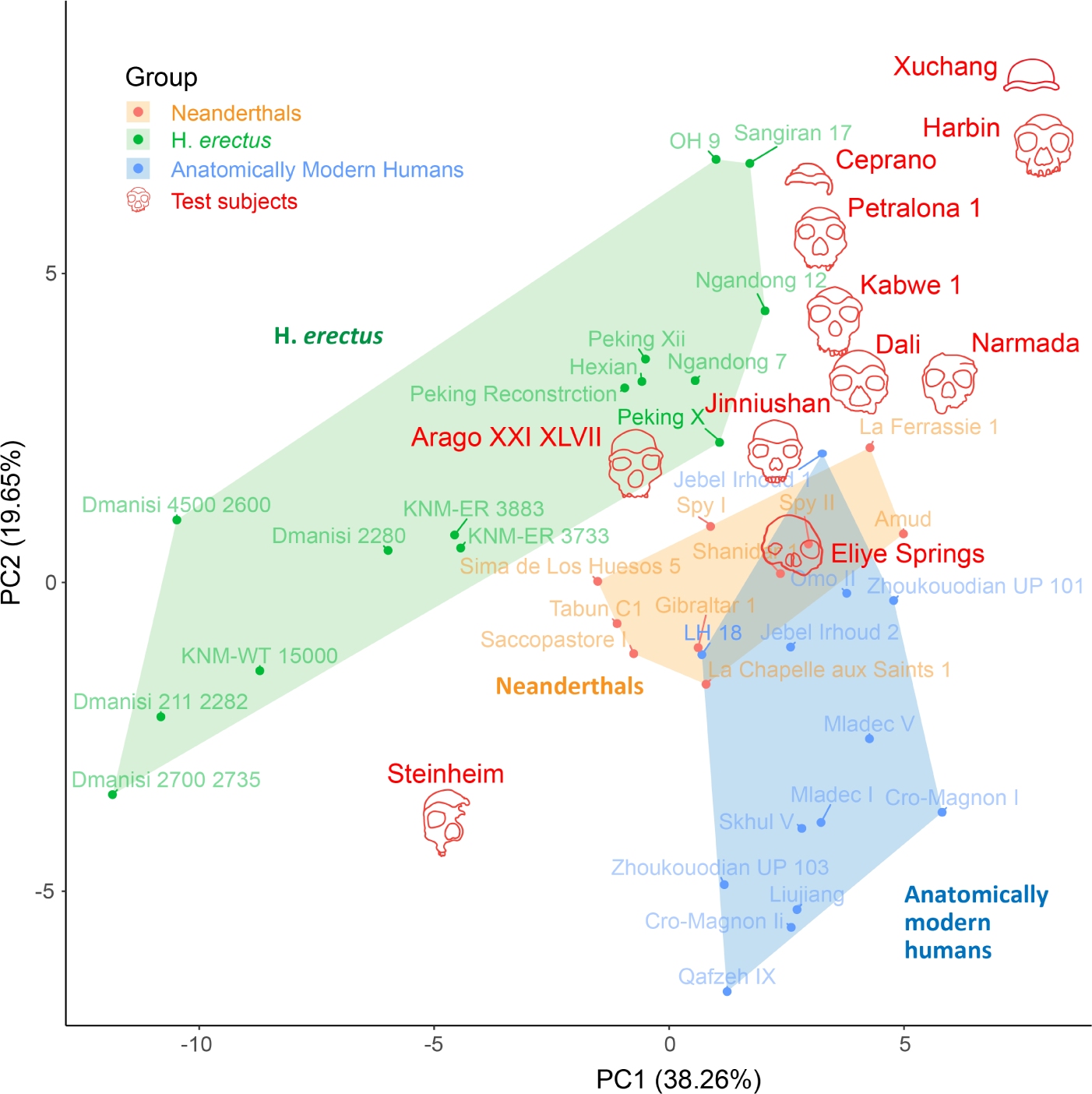
Principal component analysis (PCA) of the specimens included in the study, based on 50 available cranial measures. The polygons draw the convex hull of the three reference groups. High-scoring test subjects cluster together, suggesting morphological similarity between them.

## 5 Discussion

Identifying Denisovans in the fossil record is a key step in understanding their evolution. However, this is a highly challenging task, as attested by the scarcity of confirmed Denisovan specimens. Here, we propose that reconstructing Denisovan morphology using gene activity patterns provides a promising way to link genetics and anatomy, and opens the window to systematically scanning debated specimens and quantifying their degree of resemblance to predicted Denisovan anatomy.

We highlighted three specimens that show a particularly high resemblance to the predicted Denisovan profile [34]: *Harbin*, *Dali*, and *Kabwe 1*. Two of these, *Harbin* and *Dali*, have previously been suggested to share links with Denisovans, either through circumstantial evidence (e.g., their temporal and geographical context) [29, 1]. *Harbin* was also associated with Denisovans based on two morphological similarities to Denisovans, specifically the exceptionally large molars and M3 agenesis [21, 27]. Here, we provided the first comprehensive genetics-based evidence of their potential link to Denisovans. More generally, we found that high-scoring specimens tend to be located in East Asia, part of the putative Denisovan habitat.

The resemblance of several specimens attributed to H. *heidelbergensis* to the Denisovan profile, and specifically the African specimen *Kabwe 1*, is particularly intriguing. The similarity between *Kabwe 1* and other high-scoring specimens is evident in all PCAs (Figure 4 and Supplementary Figure 10), suggesting that this resemblance is not limited to Denisovan-like phenotypes. Despite our limited knowledge of the true range of Denisovans, it is unlikely that they reached Southern Africa, hence *Kabwe 1* is unlikely to be directly positioned on the Denisovan lineage. A more plausible explanation is that the resemblance of some of these specimens to Denisovans reflects a proximal phylogenetic affinity with Denisovans. For example, H. *heidelbergensis* specimens were positioned either close to the split between modern and archaic humans or close to the split between Neanderthals and Denisovans [29]. If H. *heidelbergensis* is indeed phylogenetically closer to the Neanderthal-Denisovan split than Neanderthals are, it is expected to exhibit an even greater similarity to the Denisovan profile than Neanderthals do. Another alternative explanation is that Denisovans might have retained several ancestral phenotypes observed in H. *heidelbergensis* [59], or that these phenotypes were affected by gene flow into Denisovans, originating from contemporaneous H. *heidelbergensis* [17].

Almost all AMH and H. *erectus* specimens in our analysis received relatively low scores, indicating that our method is not universally permissive, and that lineages that are relatively far from Denisovans do not tend to align with the profile. Neanderthals, which are expected to share many phenotypes with Denisovans (based both on their phylogeny as well as supported by our reconstructed profile), tend to show higher scores. However, even within Neanderthals, only two specimens out of 15 received scores comparable to *Harbin*, *Dali*, and *Kabwe 1* (Fig. 11). Overall, the rare occurrence of false positives (high-scoring non-Denisovan specimens), the high accuracy (*>*85%) of the anatomical profiling [34], and the predictions that were later observed in confirmed Denisovan specimens (Specifically in *Xiahe 1* [10]), support the robustness of this method in detecting Denisovan-like morphology.

The Denisovan morphological reconstruction [34] is based on a comparative analysis between AMHs, Neanderthals, and a single Denisovan specimen. We acknowledge that the morphology of a single individual cannot encompass the spatio-temporal morphological variation of an entire population. This is especially relevant in the case of Denisovans, who had a complex population structure, with deeply divergent lineages separated as early as 350 Kya [60]. This divergence, along with the wide spatial distribution of Denisovans in Asia, likely resulted in morphological variability within Denisovans [27, 1]. As a result, our reconstruction approach is likely to miss some non-fixed derived Denisovan phenotypes. However, most predictions in the original reconstruction, as well as in this work, stem from either AMH-derived or archaic-derived changes, suggesting that they are synapomorphic in Denisovans. In fact, only two measures used in this work are strictly based on Denisovan-derived phenotypes (palate length and biparietal breadth). Therefore, the morphological profile tested in this work likely consists of traits that are shared by most Denisovans.

Our work showcases the potential of gene regulatory information as a powerful predictive tool for inferring complex phenotypes. By demonstrating the robustness of this approach, we envision that similar methodologies will become instrumental in inferring the phenotypes of other extinct groups. As our understanding of gene regulation deepens, and as insights into ancient gene regulation in non-skeletal tissues emerge [30, 61, 62, 32, 33, 63], we anticipate that these techniques will expand beyond the skeletal system to include tissues that are not accessible through traditional methods.

## 6 Acknowledgements

D.G. was supported by the Center for New Scientists at the Weizmann Institute of Science and the Kahn Family Research Center for Systems Biology of the Human Cell. This study was also funded by the Israel Science Foundation ISF grant 2436/22 (to L.C). L.C. is the Snyder Granadar chair in Genetics. G.H and U.S were supported the Braginsky Center for the Interface between Science and the Humanities at the Weizmann Institute of Science. G.H was funded by the department of Physics of Complex Systems at the Weizmann Institute of Science and the Israel Academy of Science and Humanities Postodctoral Fellowship. The research was supported by the Israel Science Foundation (Grant No. 1745/21 awarded to U.S). We thank the Computational Archaeology Laboratory at the Institute of Archaeology, The Hebrew University of Jerusalem for their support. We would like to thank Chris Stringer, Erella Hovers, and Anna Belfer-Cohen for useful advice and feedback.

## 8 supplementary figures

**Supplementary Figure 1:**
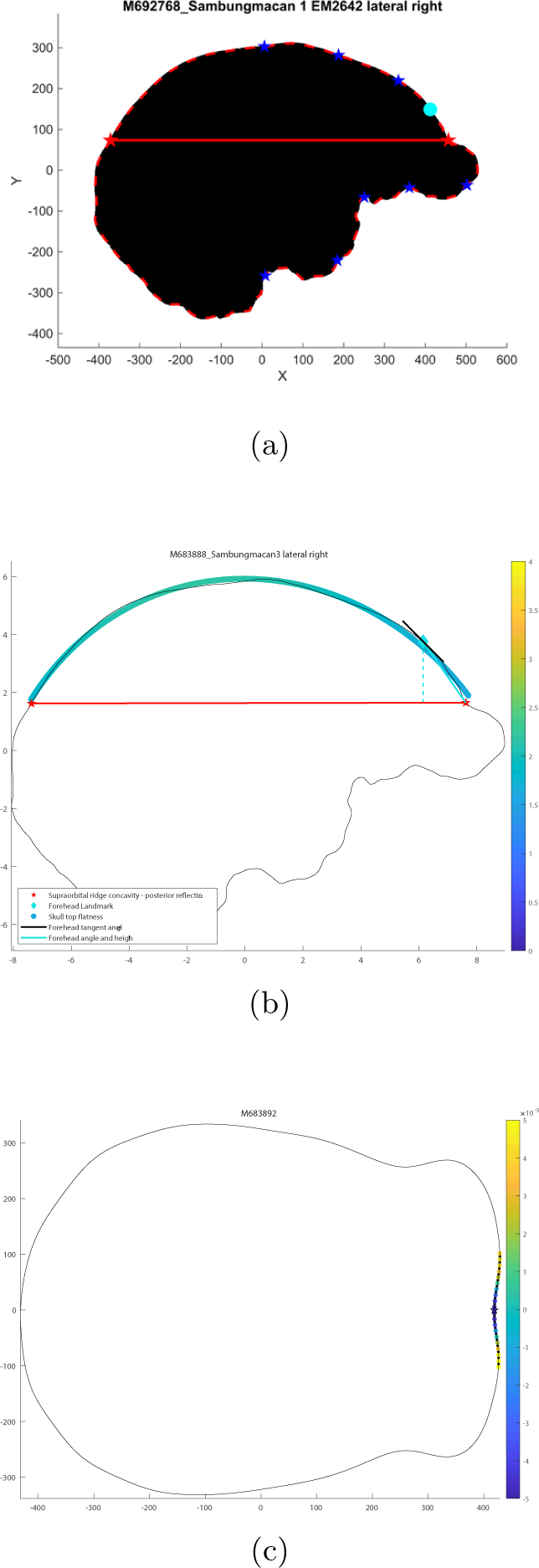
Illustration of the procedure for obtaining measurements from cranial images, with specimen Sambugmacan 1 as an example. (a) Graphical user interface for selecting critical points in the lateral view. The calvarial region is defined by the outline curve above the segment connecting the critical point (right red star) to its posterior reflection (left red star). The cyan circle indicates the position of the48forehead point. (b) Cranial top flatness values displayed using a color gradient, with the forehead height marked by a dashed line. (c) Curvature of the glabellar region in the superior view, with the glabella position indicated by a star.

**Supplementary Figure 2:**
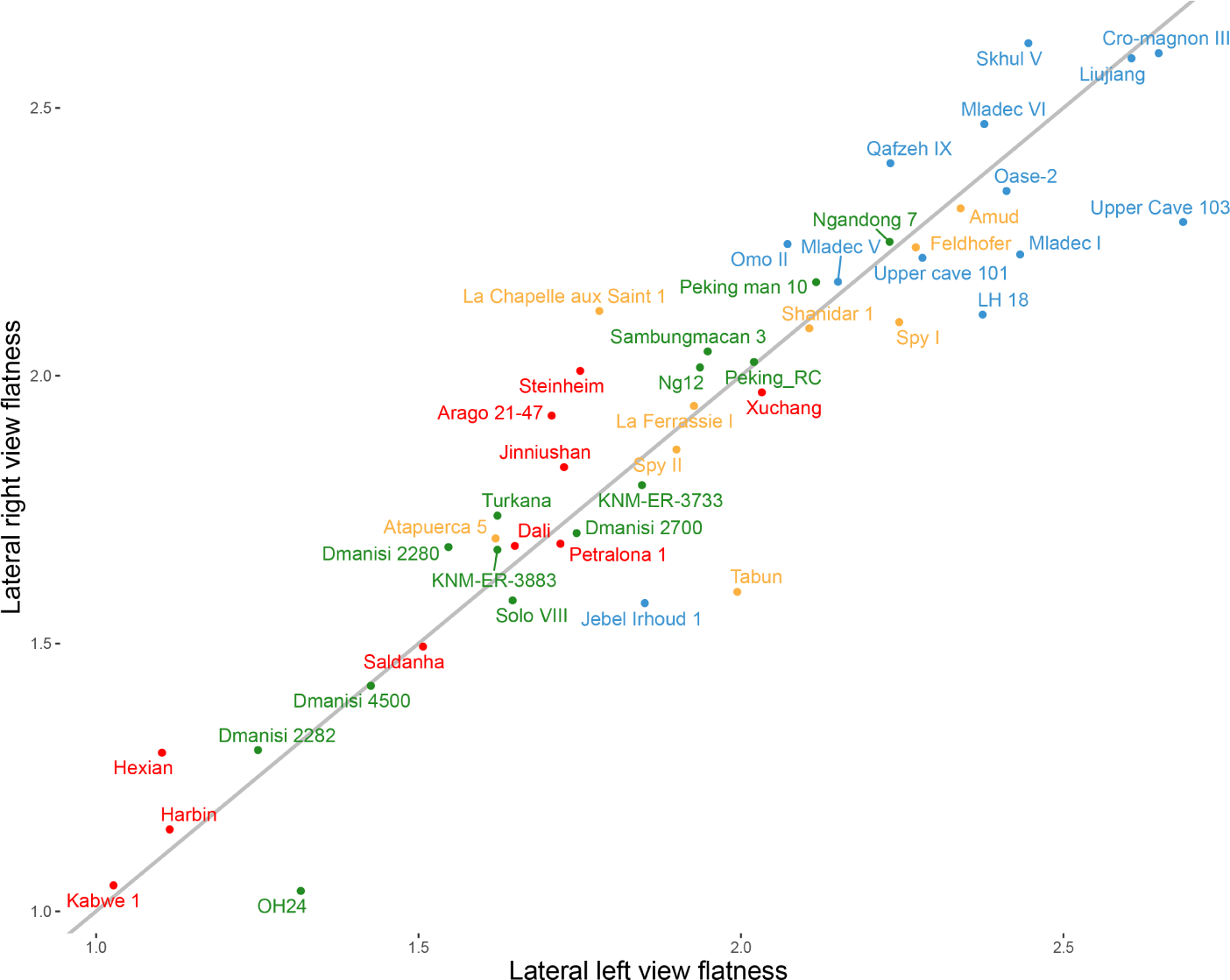
Comparison of calvarial curvature values estimated by the left and right views of the crania (*R* = 0.94). Only specimens with intact contour of the calvarium are presented (blue = AMHs, yellow = Neanderthals, green = H. *erectus*, red = Middle Pleistocene specimens). Gray line depicts the curve *y* = *x*.

**Supplementary Figure 3:**
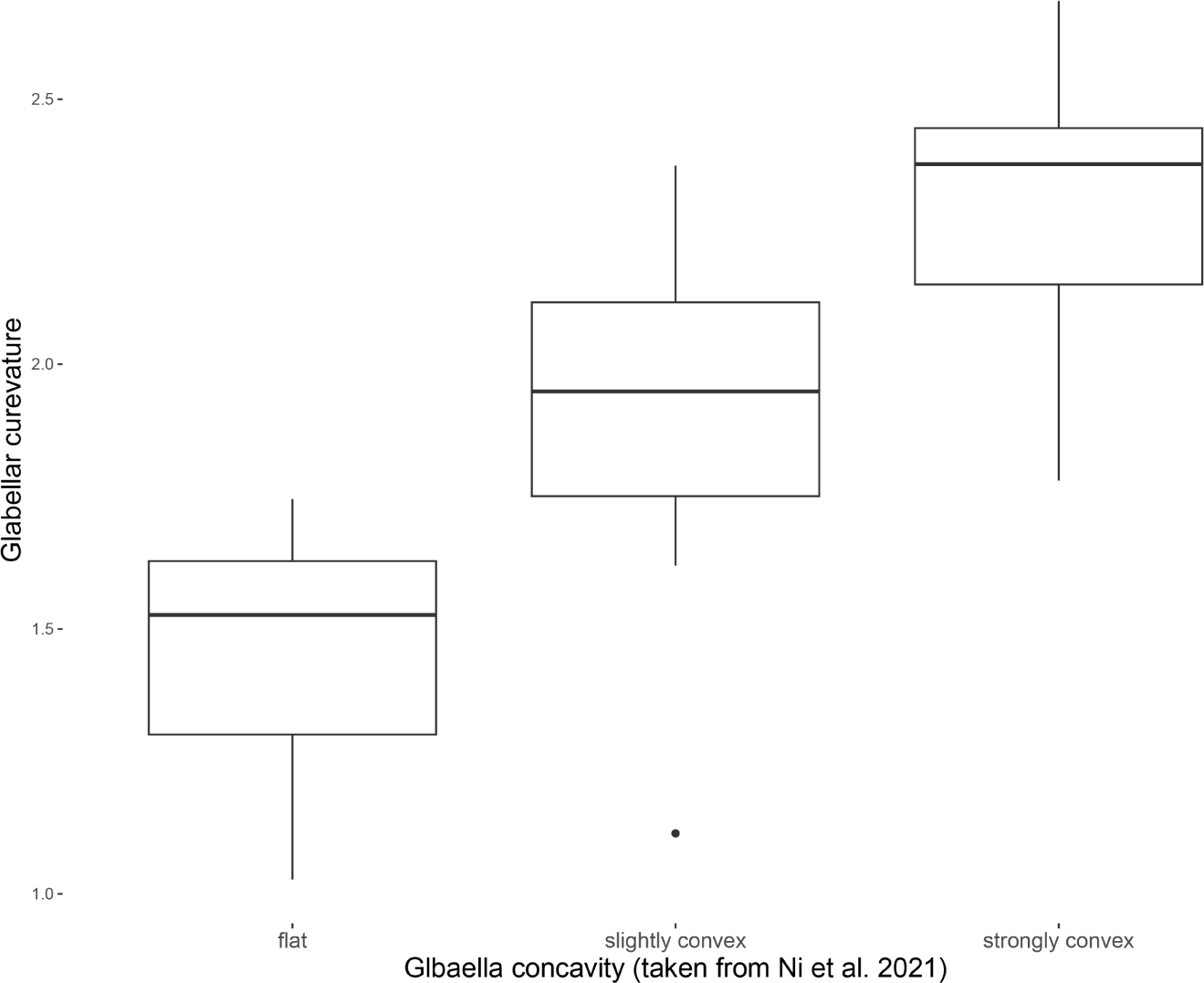
Distribution of our calvarial curvature values within each of the previously described discrete classifications of Ni *et al.*[21]

**Supplementary Figure 4:**
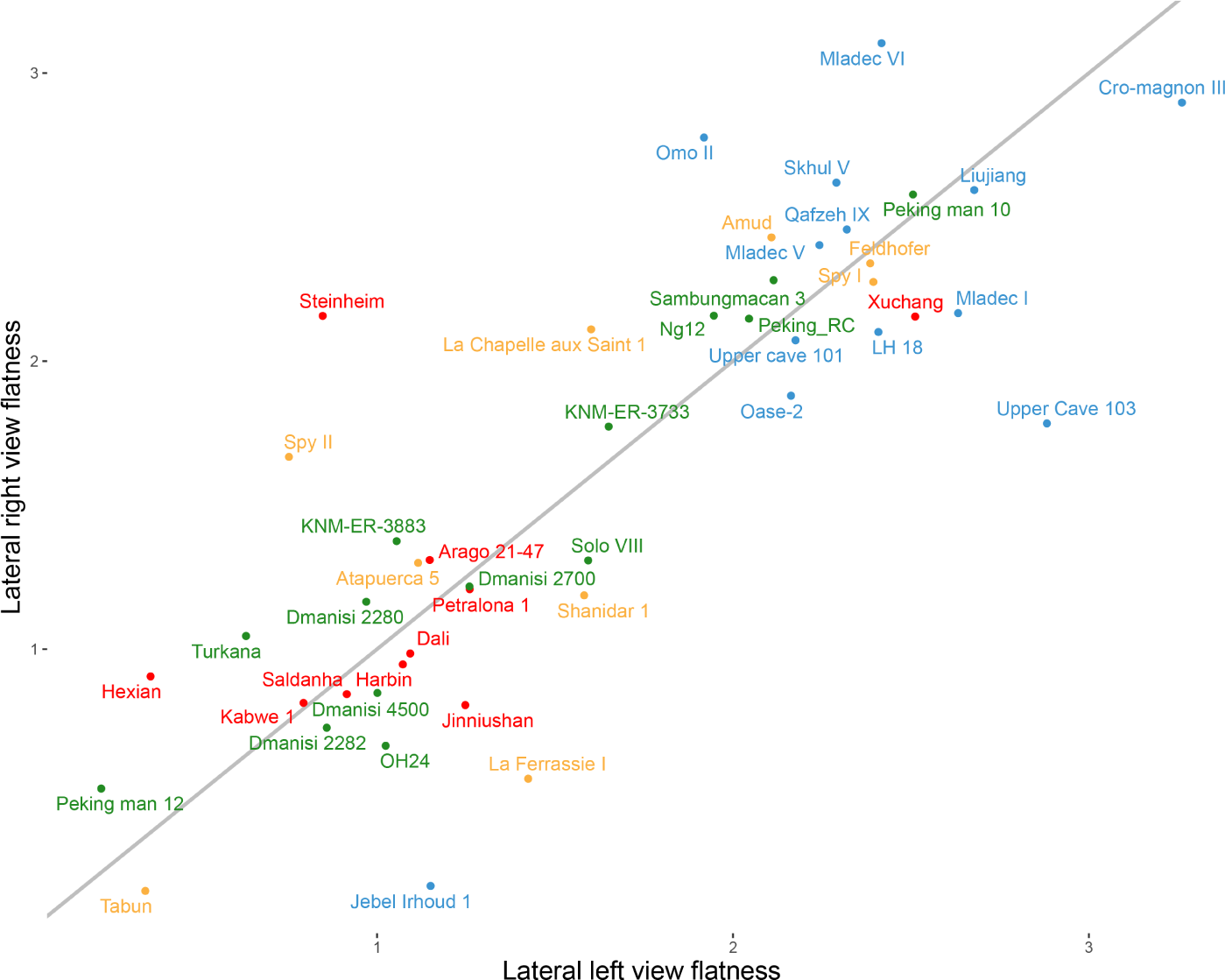
Comparison of forehead height estimated by the left and right views of the crania (*R* = 0.57). Only specimens with intact contour of the calvarium are presented (blue = AMHs, yellow = Neanderthals, green = H. *erectus*, red = Middle Pleistocene specimens). Gray line depicts the curve *y* = *x*.

**Supplementary Figure 5:**
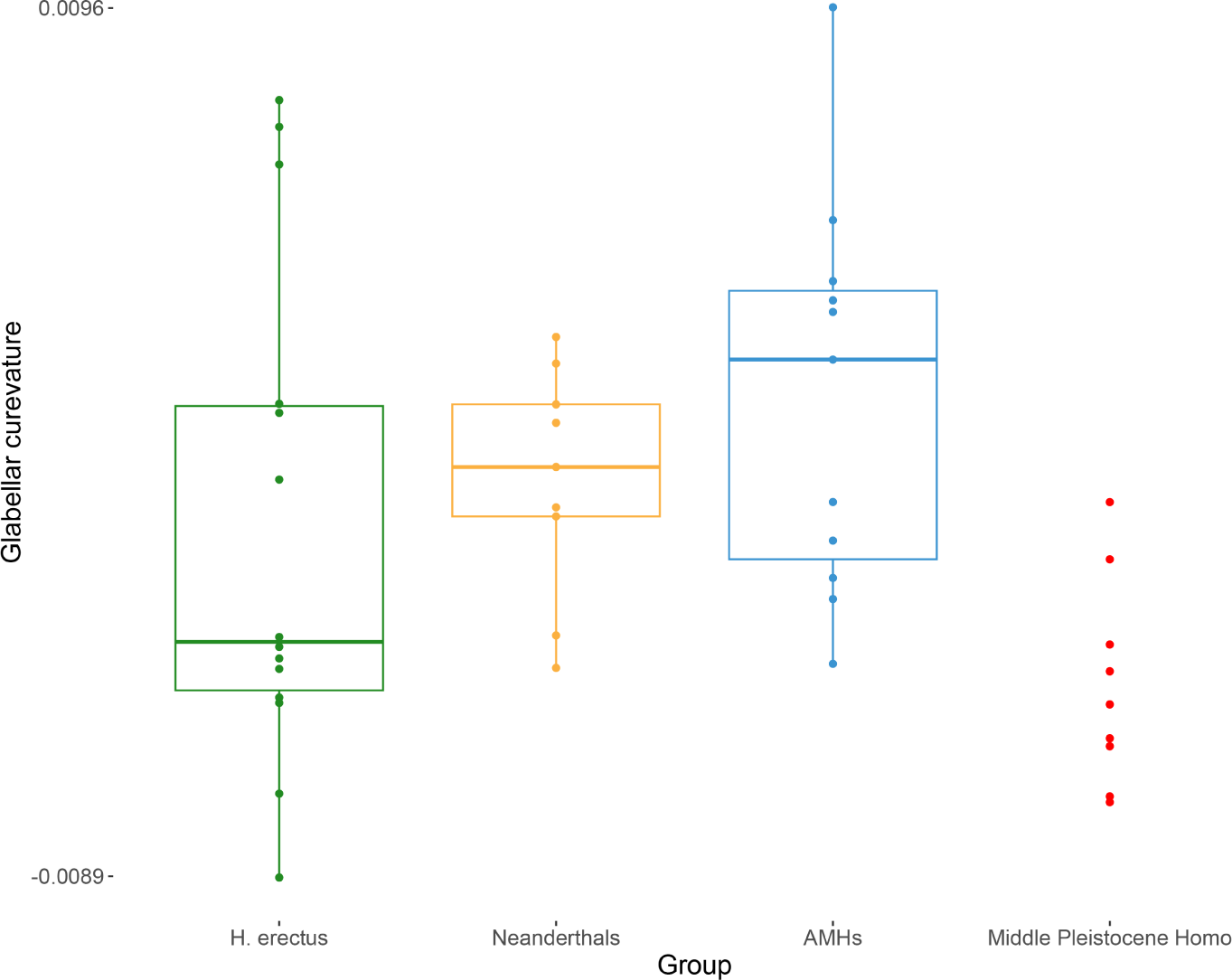
Distributions of our glabellar curvature values within each of the human lineages, as well as their value in the test subjects.

**Supplementary Figure 6:**
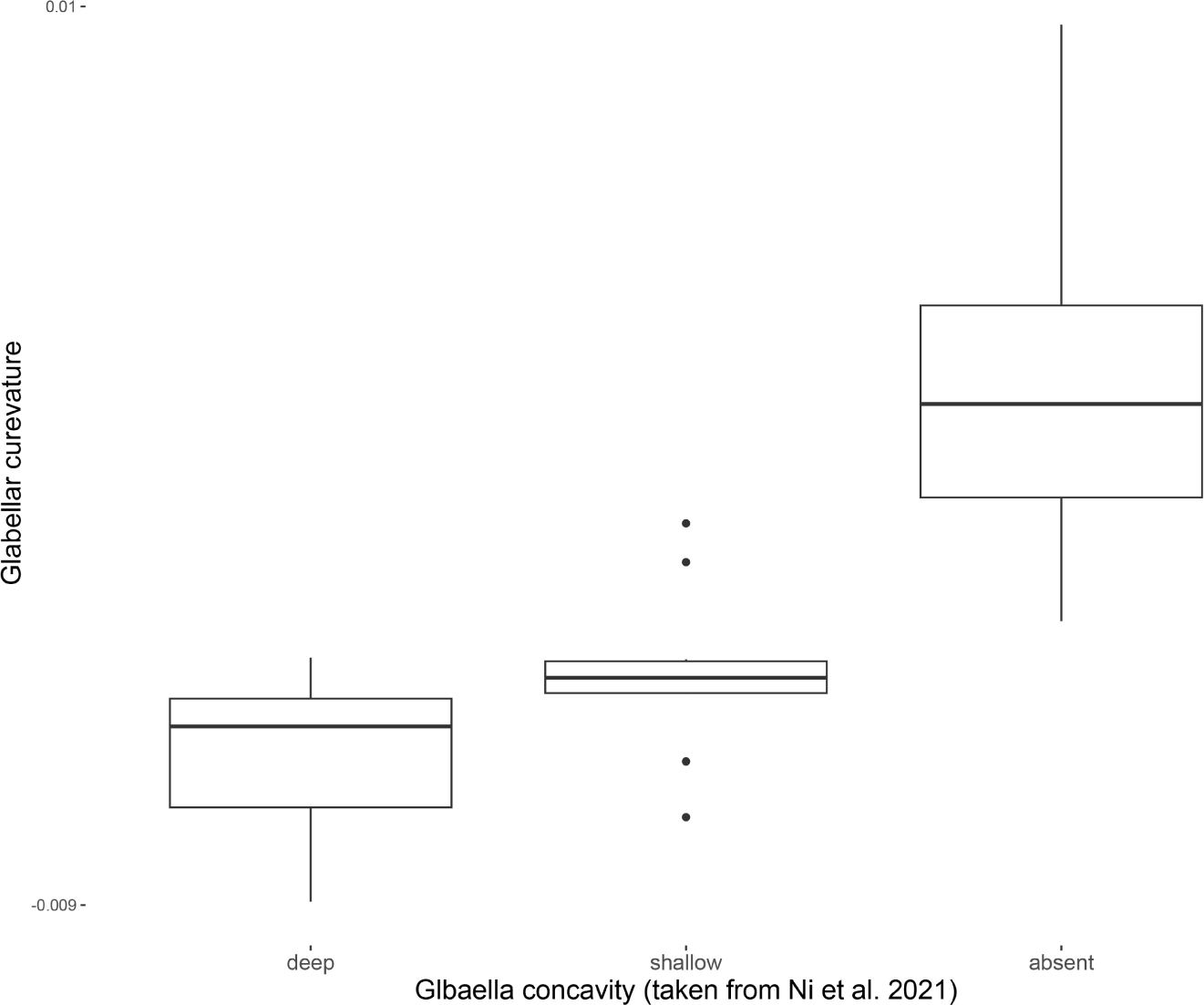
Distribution of our glabellar curvature values within each of the previously described discrete classifications of Ni *et al.* [21]

**Supplementary Figure 7:**
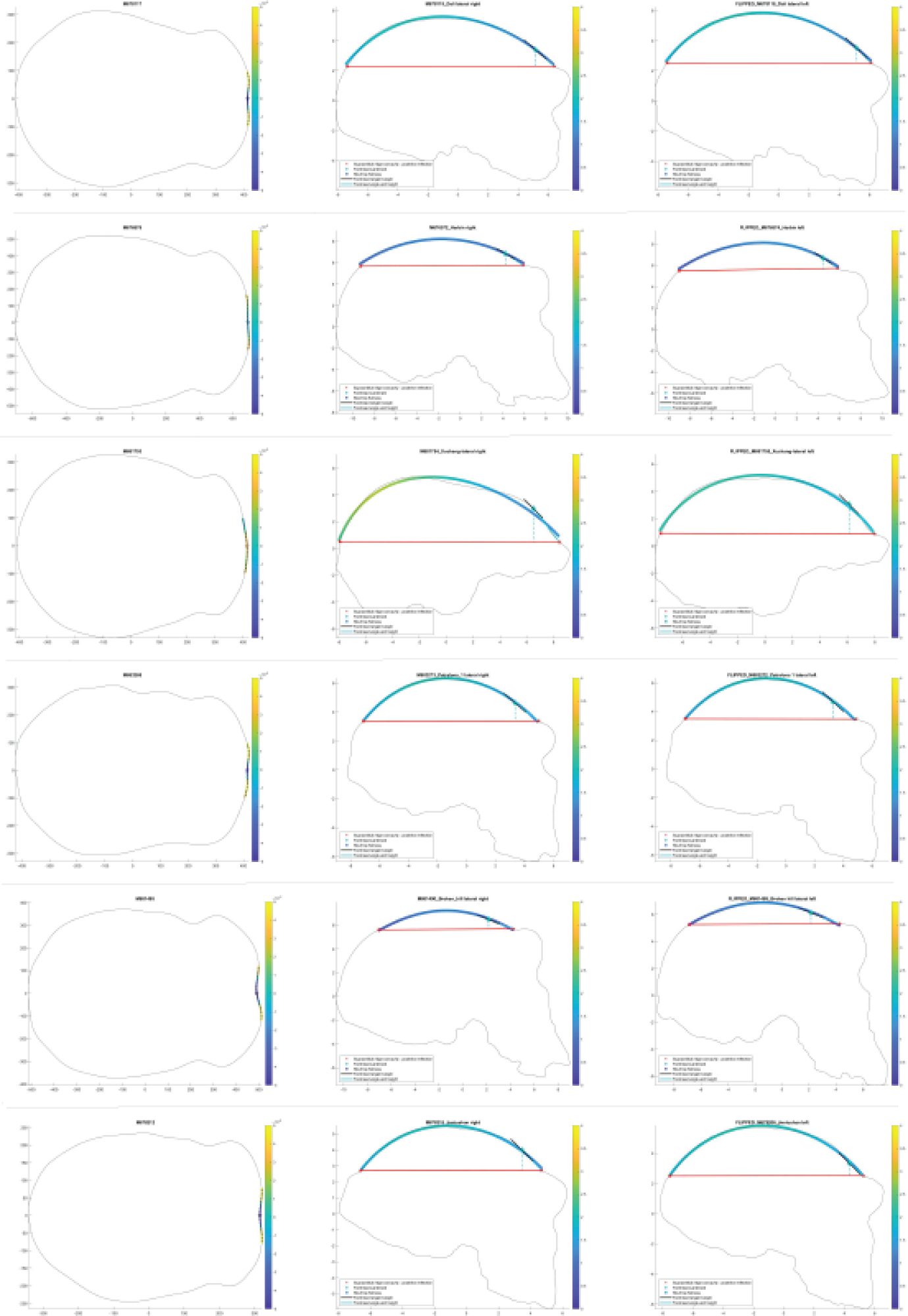
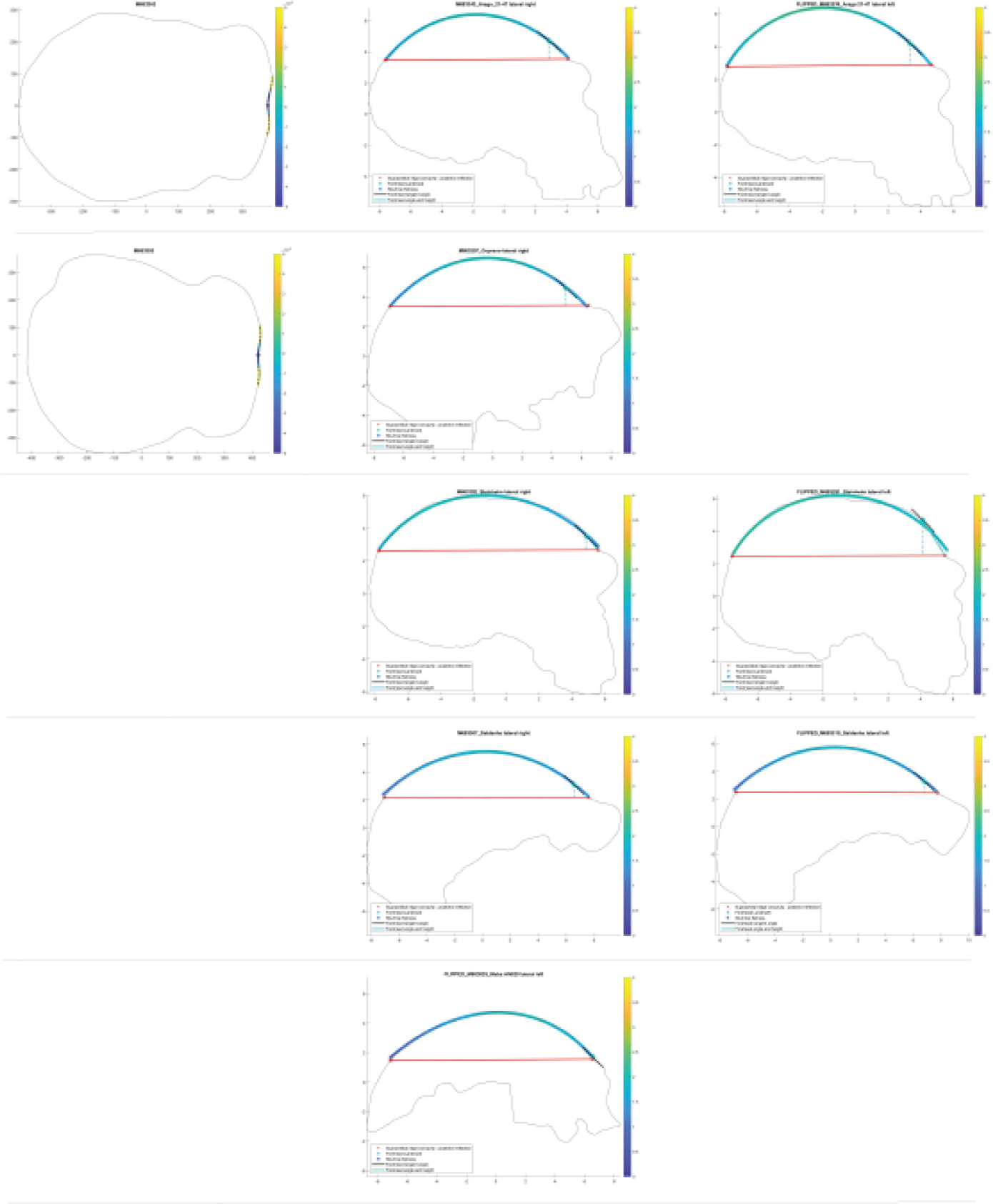
Curves used to calculate calvarial curvature, forehead height, and glabellar curvature for the test subjects. From left to right: superior view, lateral right view, and horizontally flipped left lateral view. From top to bottom: *Dali*, *Harbin*, *Xuchang 1*, *Petralona 1*, *Kabwe 1*, *Jinniushan*. Curves used to calculate calvarial curvature, forehead height, and glabellar curvature for the test subjects. From left to right: superior view, lateral right view, and horizontally flipped left lateral view. From top to bottom: *Arago 21-47*, *Ceprano*, *Steinheim*, *Saldanha*, *Maba*. Curves that were not used in the final analysis due to fragmentation or distortion are not shown.

**Supplementary Figure 8:**
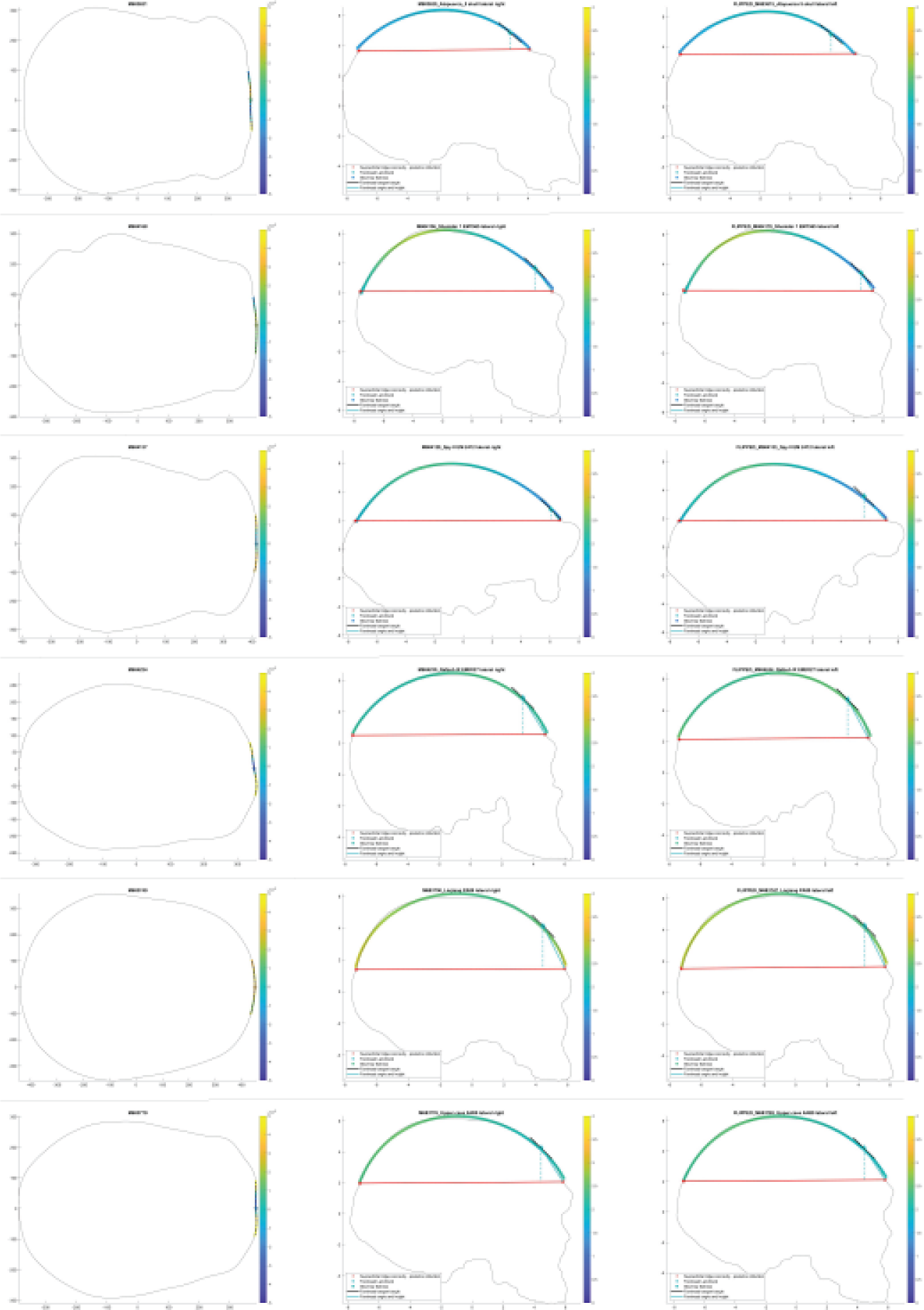
Curves used to calculate calvarial curvature, forehead height, and glabellar curvature for some of the Neanderthal and AMH specimens. From left to right: superior view, lateral right view, and horizontally flipped left lateral view. From top to bottom: *Sima de los Huesos 5*, *Shanidar 1*, *Spy II*, *Qafzeh IX*, *Liujiang*, *ZKD Upper cave 101*.

**Supplementary Figure 9:**
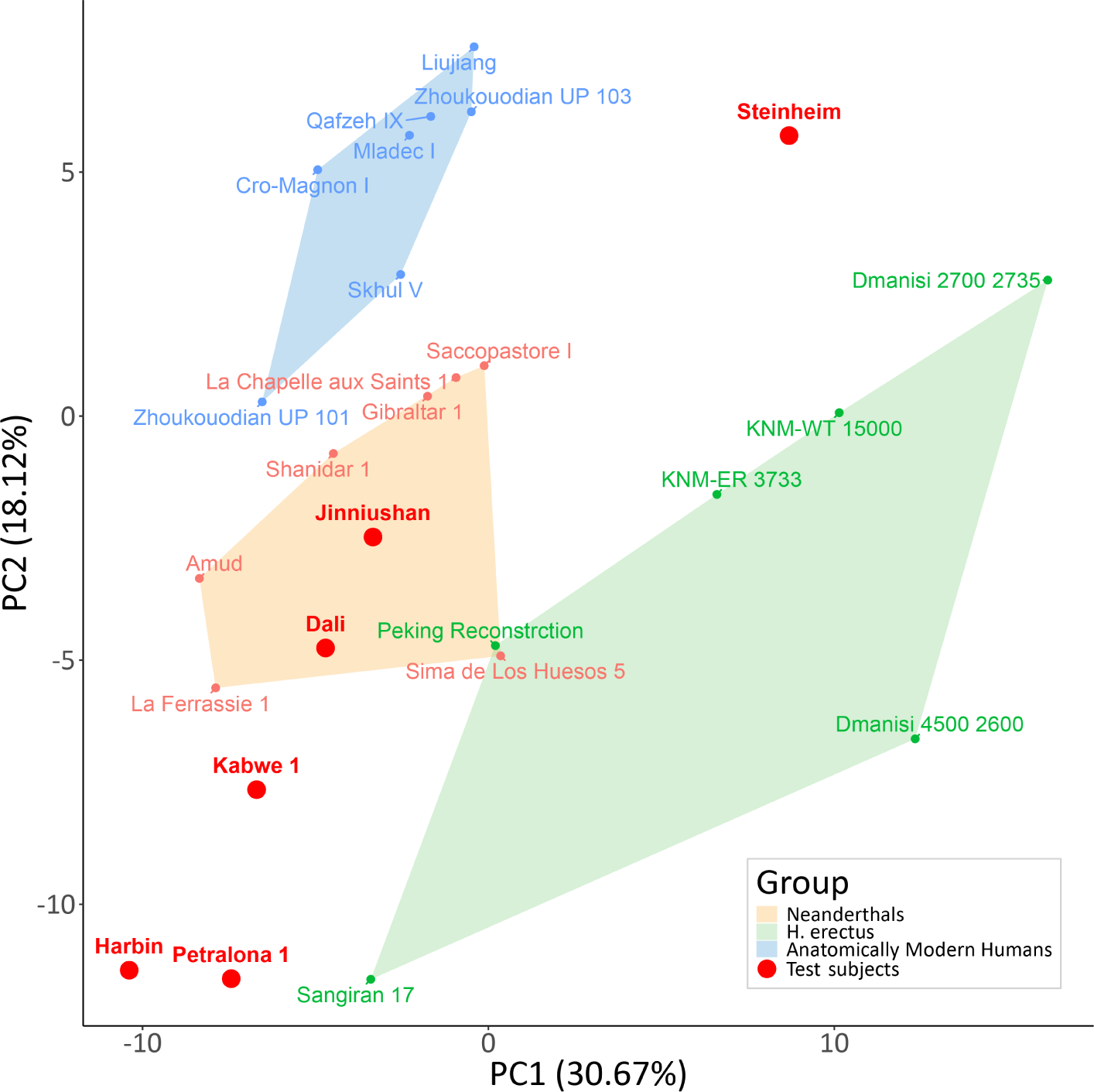
PCA of the specimens included in the study, based on 139 available cranial measures. The polygons draw the convex hull of the three reference groups. The filtering steps in this analysis are flipped compared to the main PCA; First, specimens with more than 15% missing measures were filtered out, and only then measures with more than 20% missing data were filtered. This order of steps enabled the inclusion of more measurements, but left out more specimens. Similarly to the main PCA, most test subjects are positioned outside the reference group clusters. However, here, Dali is positioned within the convex hull of Neanderthals.

**Supplementary Figure 10:**
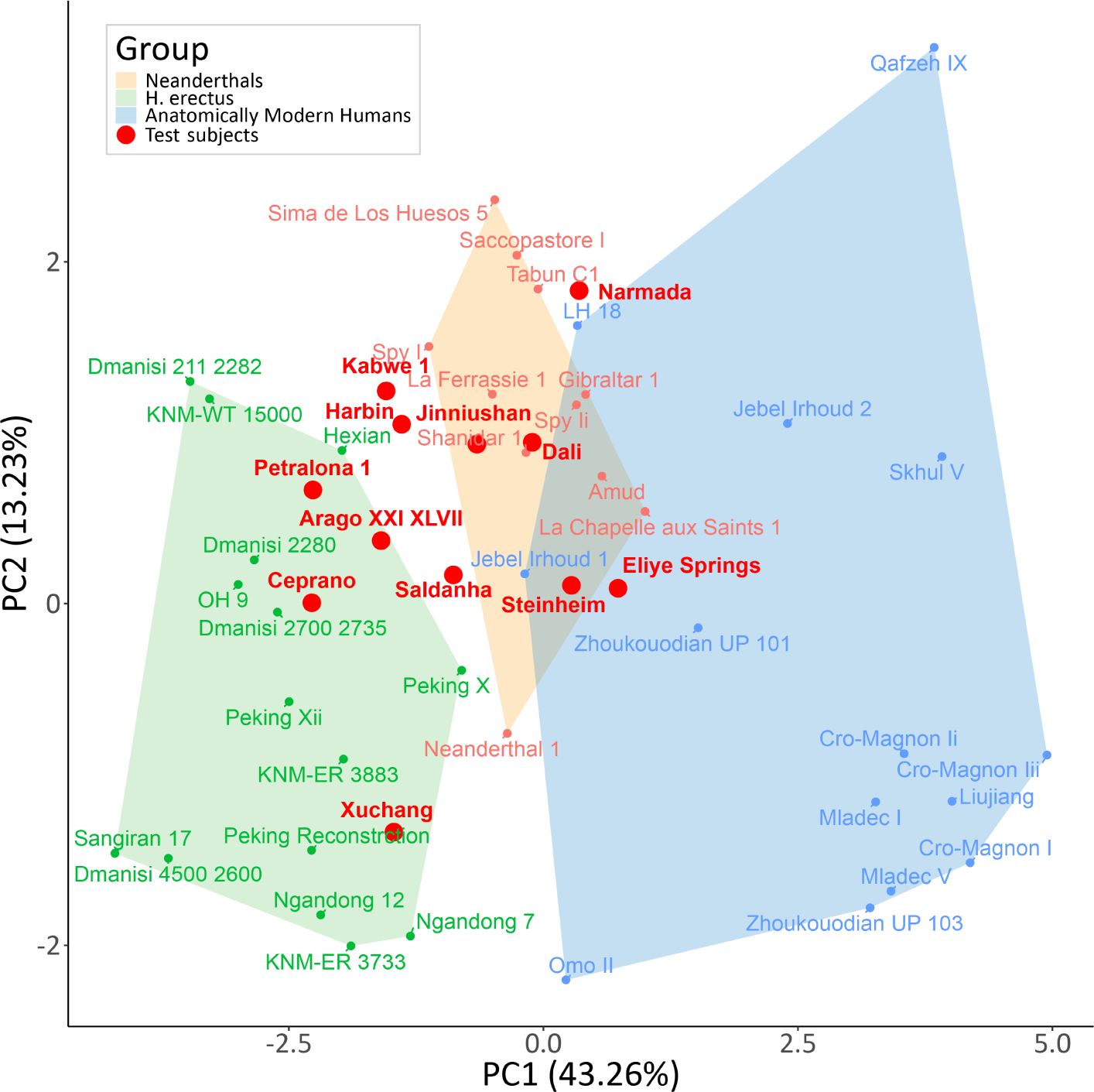
PCA of the specimens included in the study, based on 13 available non-metric cranial measures. The polygons draw the convex hull of the three reference groups. In this analysis, we excluded all metric measures, leaving only angles and ratios, after applying the filtering described for the main PCA. In this PCA, the reference groups remain mostly separated. However, most test subjects in this analysis fall within the convex hulls of the reference groups.

**Supplementary Figure 11:**
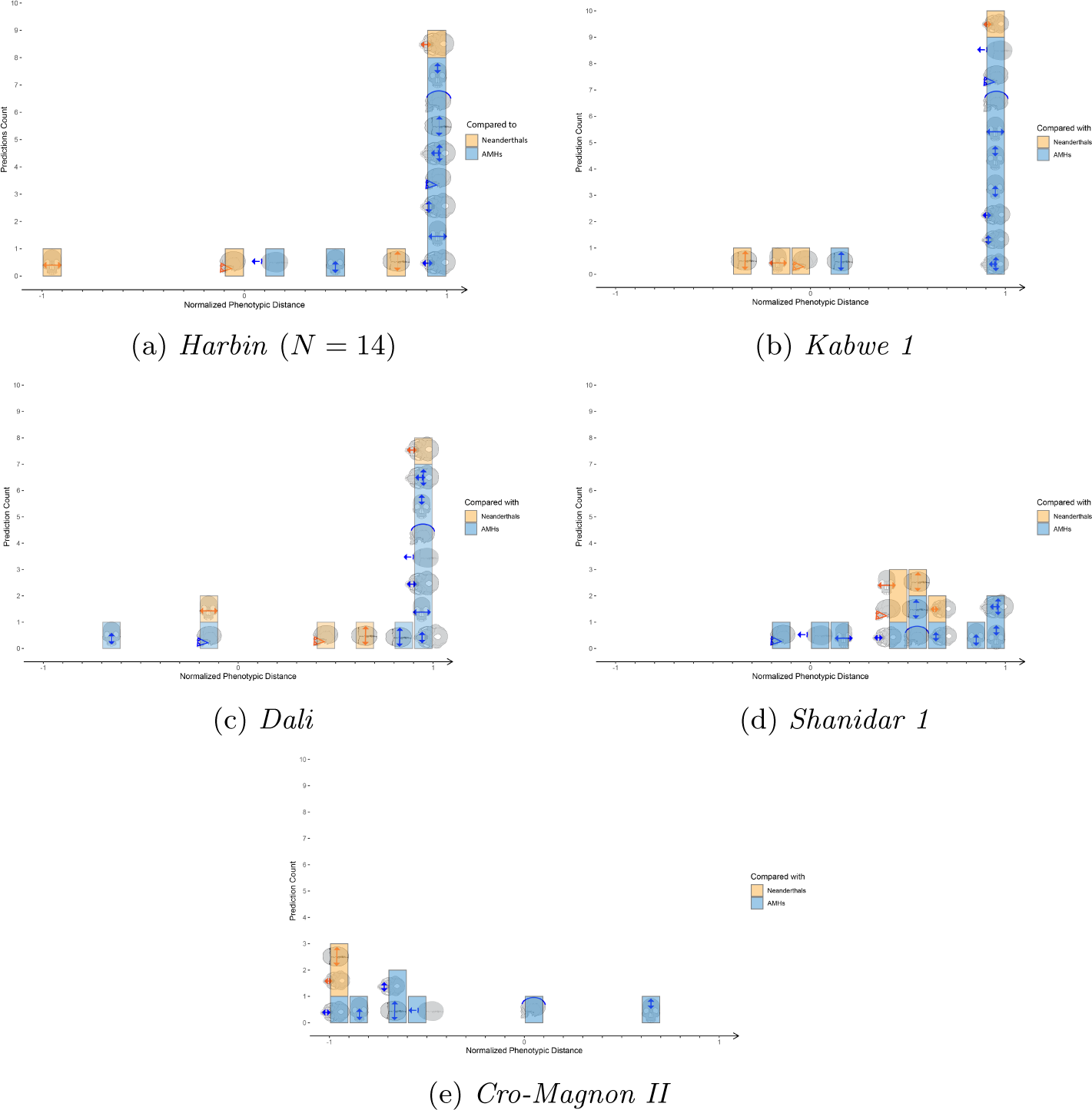
Overall match of selected specimens to the Denisovan profile, represented as stacked histograms of phenotypic distances. Blue bars stand for measurments that are compared to AMHs, yellow bars stand for measurements that are compared to Neanderthals. (a) For *Harbin*, 12 out of 14 predictions align with the Denisovan profile. (b) For *Kabwe 1*, 11 out of 14 predictions align with the Denisovan profile. (c) For *Dali*, 11 predictions align with the Denisovan profile. (d) For *Shanidar 1*, 13 out of 14 predictions align with the Denisovan profile. (e) For *Cro-Magnon II*, 2 out of 9 predictions align with the Denisovan profile.

